# The Computational and Neural Basis of Zero-Shot Control in Dynamic Pursuit

**DOI:** 10.64898/2026.03.30.715455

**Authors:** Daehoon Kim, Jungsuk James Lee, Benjamin Y. Hayden, Seng Bum Michael Yoo

**Affiliations:** Department of Biomedical Engineering, Sungkyunkwan University; Department of Intelligent Precision Healthcare Convergence, Sungkyunkwan University; Department of Biomedical Engineering, Case Western Reserve University; Department of Neurosurgery, Baylor College of Medicine

**Author notes:** Equal contribution (Daehoon Kim, Jungsuk James Lee).

**Keywords:** Spotlight attention, Relational structure, Affordance, Zero-shot generalization, Change-of-mind, Dorsal anterior cingulate cortex, Nonhuman primate

## Abstract

Biological agents flexibly adapt their behavior to novel goals and environmental demands without additional training, yet the potential computational principles supporting such control remain unclear. Here, we investigate whether three cognitive constructs: relational structure, spotlight attention, and affordance computation contribute to flexible control of dynamic pursuit in novel environments. We hypothesize that these constructs support flexible generalization in an embodied dynamic pursuit task, in which successful behavior requires continuous integration of inter-entity relations, reward, and action feasibility. By implementing these constructs within a multi-module graph convolutional network, we show that the model achieves zero-shot transfer across novel pursuit scenarios without additional training. Although not explicitly trained to do so, the model also exhibits change-of-mind behavior, which is a flexible control exhibited by biological agents. Neural recordings from the primate dorsal anterior cingulate cortex revealed population-level signatures related to the computational constructs examined in our model, providing a basis for investigating their potential biological relevance.

## Introduction

The capacity for flexible control under novel environmental demands is a hallmark of intelligence^1,2^. Consider wolves hunting elk across seasons: in deep snow or open grassland, they single out a target from the herd, sustain pursuit as terrain and group structure shift from moment to moment, and disengage when conditions render capture infeasible. Such behavior requires adapting to a changing environmental structure without additional training on novel configurations. What cognitive constructs enable this form of real-time, zero-shot adaptation, and how are they implemented in biological circuits?

Dynamic pursuit is a prime example of flexible control, posing foundational computational challenges for zero-shot transfer. First, an agent must adapt when new entities appear, because introducing agents with new behavioral policies can alter pursuit dynamics^3^. Second, an agent must selectively commit to one entity among many, as encoding all entities simultaneously risks combinatorial explosion^4,5^. Third, target selection must be grounded in physical feasibility: persisting toward a high-reward target may be suboptimal if capture is unattainable^6,7^. Dynamic pursuit requires all three continuously and in real time, making it a stringent testbed for naturalistic zero-shot generalization.

Our central hypothesis is that three cognitive constructs, namely relational structure, spotlight attention, and affordance computation, contribute to flexible control of dynamic pursuit in novel environments. The explicit encoding of relational structure enables abstraction across entities with distinct goals and behavioral policies — such as prey versus predator — thereby allowing rapid behavioral adaptation^8–10^. Spotlight attention prevents a combinatorial explosion as the number of entities grows, allowing stable commitment^11–14^. Affordance computation grounds target selection based on physical feasibility, enabling the agent to disengage regardless of reward magnitude^15–17^. We test this hypothesis through targeted manipulations of affordance computation, selective attention, and relational structure, and evaluate their contribution to generalization.

Despite advances in computational models capable of zero-shot transfer in domains such as physical and conceptual navigation^9,18,19^ and categorization^20,21^, zero-shot generalization in dynamic, embodied pursuit remains largely unexplored. In these prior domains, generalization has been linked to hippocampal relational abstraction^22,23^ or structured priors in parietal cortex^24^. However, embodied and interactive pursuit imposes a qualitatively different demand: generalization must unfold through continuous monitoring and real-time control of action under evolving physical feasibility constraints. This requirement implicates neural mechanisms that not only represent static information from the current configuration but also dynamically arbitrate commitment, monitor conflict, and select actions as the configuration evolves. Given these additional demands, the dorsal anterior cingulate cortex (dACC) is a candidate region to support this form of control. dACC has been implicated in conflict monitoring^25,26^, evidence accumulation^27^, and contextual adaptation during decision-making^28–30^, functions that are well suited to the demands of pursuit. We therefore examined whether dACC population activity reflects the computational operations instantiated in each module of our architecture.

## Results

### Behavioral Similarity Between Computational Model and Subjects

To test the contribution of the three cognitive constructs for flexible control, we used a dynamic prey-pursuit task^31,32^, in which a subject navigates a circular avatar using a joystick to pursue a square-shaped prey item with varying speeds and reward values. In the biological experiments, the task consisted of two conditions: in task 1, the subject pursued a single prey (Fig. 1a, top), whereas in task 2, it pursued two prey (Fig. 1a, bottom). The trial goal and termination condition (capture of one prey within 20 seconds) were identical across both conditions (task 1: n = 5,847; task 2: n = 6,022). We translated this task into a training environment for artificial agents. The model was trained only under the task 1 condition, with slower prey speeds and a smaller arena (see *Training environment for agent* in Methods), to learn the task structure. As a result, all subsequent conditions, including multi-prey scenarios and altered environmental dynamics (see Generalization Protocols and Evaluation Metrics in Methods), constituted genuine zero-shot tests that required the model to generalize without any additional parameter updates.

**Fig. 1.**
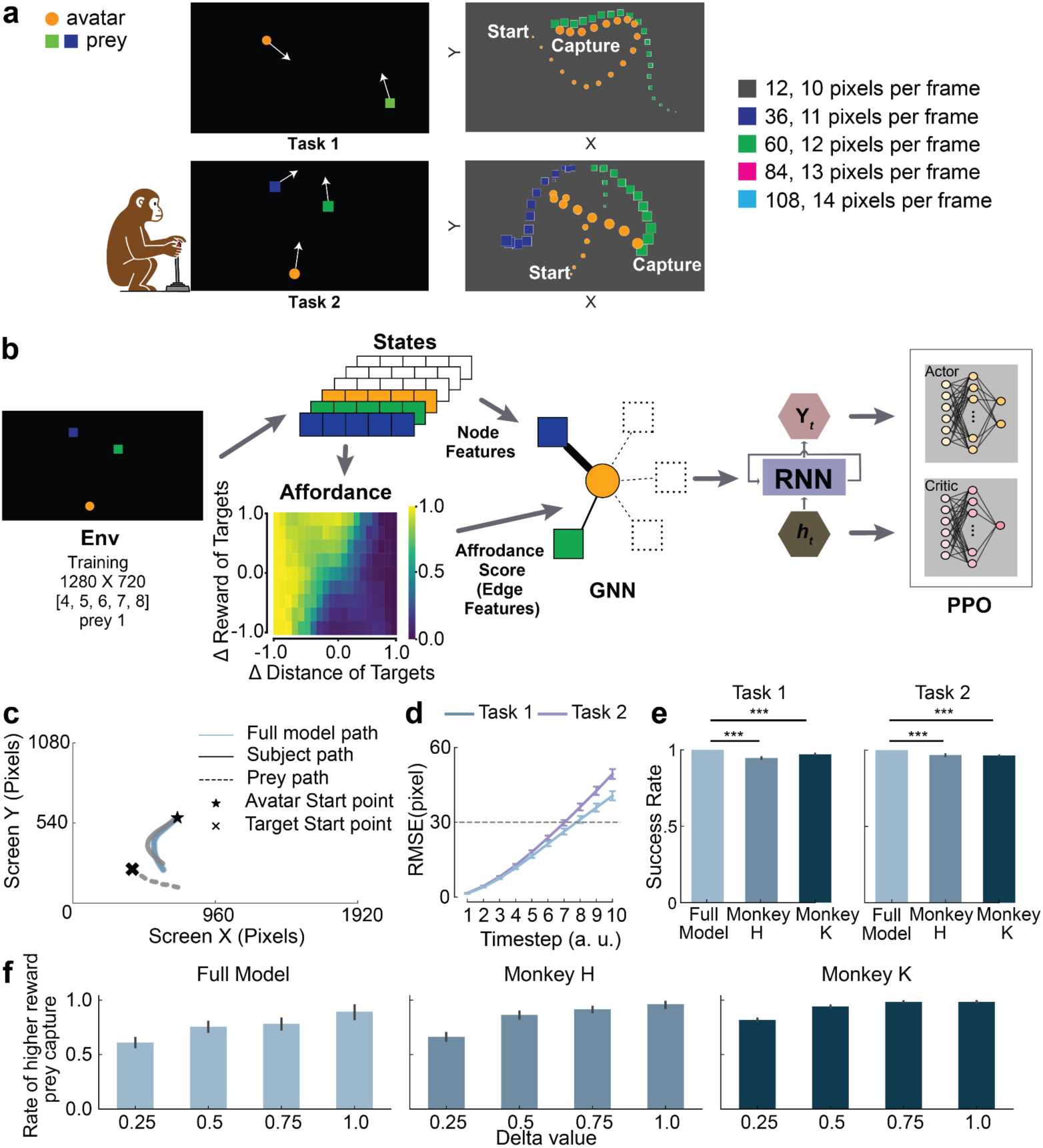
Task Description and Model Architecture. **a)** Task Schematics. The subject uses a joystick to control the circular avatar to pursue prey, denoted as a square avatar on a computer screen. The top of the screen shows one prey (task 1), while the bottom of the screen shows two prey (task 2). Box colors match the prey colors in the task and denote reward magnitude (a.u.) and maximum movement speed (pixels per frame). b) Model Architecture. The model consists of four modules: (1) Affordance module; (2) Graph Convolutional Network (GCN); (3) Recurrent Neural Network (RNN); and (4) PPO Actor-Critic module. c) Example trajectories and corresponding fit trajectories generated by the model. d) Root Mean Squared Error (RMSE) of the predicted rollout trajectory for the two tasks. The dashed line indicates the avatar’s radius (30 pixels). Error bars are the standard error of the mean (s.e.m). e) Success rate comparison of Task 1 (1-prey; left) and Task 2 (2-prey; right) for the model agent (skyblue), Subject H (blue), and Subject K (dark blue). f) Success rate of intercepting the higher-reward prey for the agent and subject based on the reward gap of two prey, denoted as “delta value.”

The model consists of four modularized components, each reflecting a distinct cognitive construct of a biological agent. First, inspired by embodied accounts of action selection, the network computes an unsupervised learned affordance score from the state representation of the avatar and prey, including their positions, velocities, and reward values^33^. Second, drawing on relational accounts of the cognitive map, a graph convolutional network (GCN) encodes entity representations through a masked softmax over learned affordance scores and applies them to each edge representing separate prey^34,35^. These edge weights implement attentional allocation, with spotlight attention corresponding to a distribution concentrated on a particular target. Third, a recurrent neural network (RNN) integrates successive graph representations, allowing action selection to depend on temporal context as well as the current input^36^. Finally, an actor-critic module maps the recurrent representation to an action distribution and a value estimate, while proximal policy optimization (PPO) trains the network parameters using reward feedback (Fig. 1b)^37^.

To assess whether the model can reproduce pursuit behaviors of biological agents, we compared the rollout trajectories from matched initial states between the model and subjects in both 1-prey (Task 1, 1,496 trials) and 2-prey (Task 2, 1,485 trials) conditions under matched environmental settings (Fig. 1c). The trajectory difference, estimated by root mean square error (RMSE), remained near the agent’s physical radius (30 pixels) for up to 8 timesteps (Fig. 1d; median at step 7: Task 1: 26.27; Task 2: 29.79; Total: 28.03; z = –3.87; p = 5.452 × 10^−5^; one-sample Wilcoxon signed-rank test). We further evaluated behavioral similarity by comparing prey interception success rates (Fig. 1e). Although the model was trained only on Task 1, interception performance reached ceiling levels (probability of 1.0) in both Task 1 and Task 2 and exceeded that of subjects (Fig. 1e; Task 1: H = 94.7%, K= 97.1%, Full model = 100%; Δ_obs_ (observed difference between subjects and model) = 3.9%; p = 1.0 × 10^−6^; Task 2: H = 96.7%; K = 96.4%, Full model = 100%; Δ_obs_ = 3.5%; p = 1.0 × 10^−6^; unpaired permutation test). Consistent with subject behavior, as the reward gap between prey increased, the model showed a significantly stronger preference for the higher-value target (Fig. 1f; β = 1.96 ± 0.26; z = 7.52; p = 5.627 × 10^−14^; binomial generalized estimating equation (GEE)). Because any agent trained to maximize reward is expected to show this preference, this effect indicates a basic behavioral correspondence between the model and subjects, but does not by itself distinguish among candidate models or establish a shared mechanism. We therefore compare models in the following sections under conditions where their predictions diverge. Together, these results show that the model reproduces basic behavioral signatures of pursuit and generalizes to 2-prey conditions despite being trained exclusively on 1-prey scenarios.

### Explicit relational structure resolves the introduction of an agent with a new goal in flexible control

During pursuit, agents may encounter entities governed by novel behavioral policies. In such cases, flexible control requires inferring these policies and structuring relations among entities to guide appropriate responses without prior experience. To test whether structuring relations confers an advantage under such conditions, we compared two architectures that differ in how they represent inter-entity interactions: a graph-based model that explicitly structures interactions and selectively regulates information flow (Full model), and a parameter-matched multilayer perceptron that lacks explicit relational structure (No-RS model) (Fig. 2a). We evaluated both models under a modified rule introducing a triangular predator with a novel behavioral policy, testing zero-shot generalization in a larger arena with faster prey. Under this rule, trials were deemed unsuccessful if the predator reached the circular avatar before it intercepted the square prey.

**Fig. 2.**
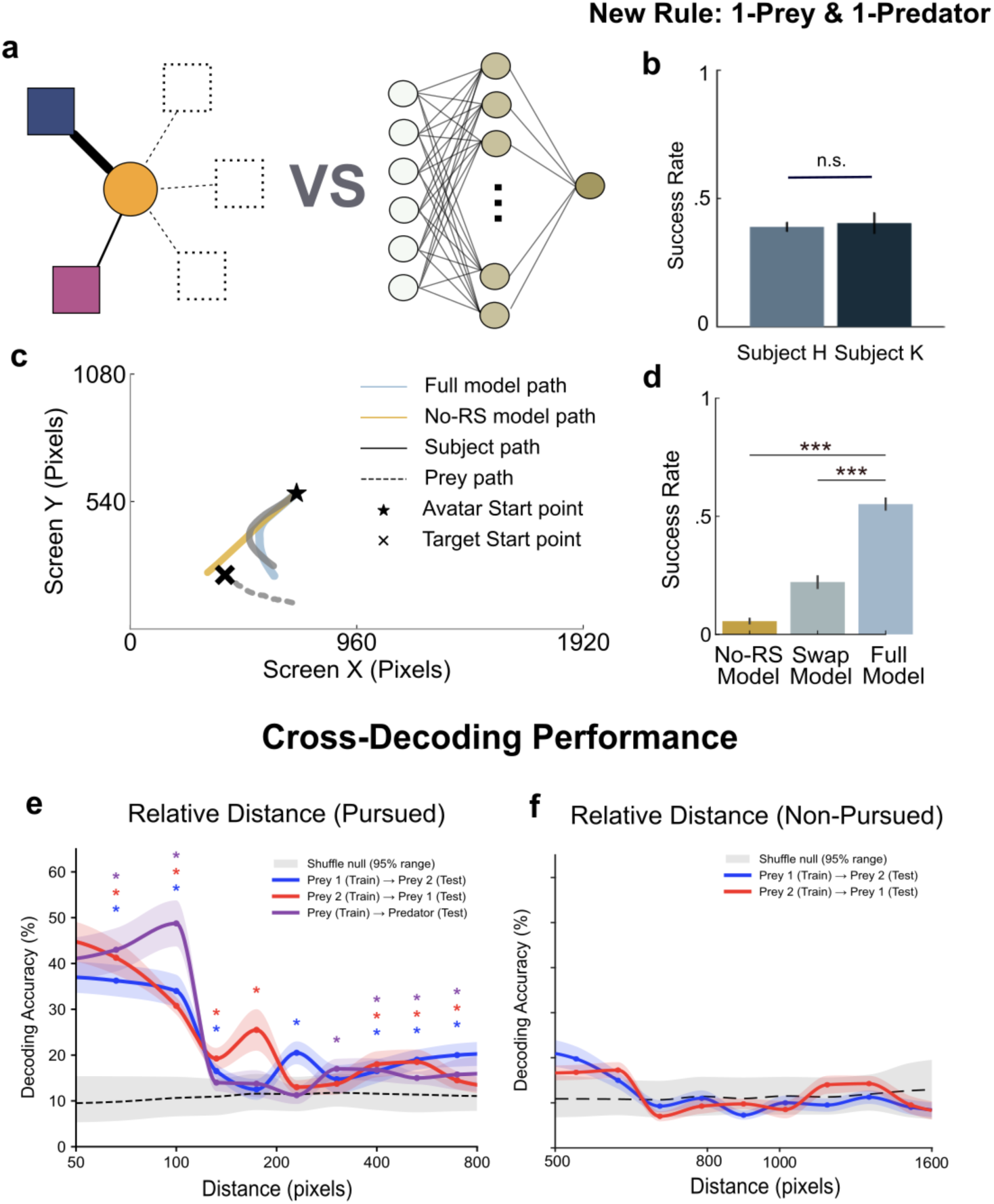
Relational Structure Enables Zero-shot Adaptation to Novel Entity Roles and is Reflected in Equivariant Population Coding in dACC. **a)** Ablation Schematics comparing the graph convolutional network (GCN), which is replaced by a parameter-matched multi-layer perceptron (MLP). b) Success rate of subject H (blue) and subject K (dark blue) of the ‘1 Prey, 1 Predator’ scenario. Error bars are the standard error of the mean (s.e.m.). c) Example trajectories and corresponding fit trajectories generated by the model. d) Success rate in the ‘1 Prey, 1 Predator’ scenario for the No-RS model in yellow, the Full models with swapped edge weights in gray-green, and the Full model in sky blue. In the swap condition, prey and predator edge weights were exchanged during evaluation without changing node features or learned parameters. e) Cross-decoding performance of relative distance (distance from self to prey) based on neural activity. The decoder is trained on one prey and tested on the other. Test results for prey 1 are shown in red, test results for prey 2 are shown in blue, and test results for the predator are shown in purple. The grey line indicates 95% of the shuffled decoders. f) The same plot as Fig. 2e but for non-pursued prey (* p < .05, ** p < .01, *** p < .001).

When the predator was introduced, subjects H and K achieved success rates of 38.97% (n = 662) and 40.44% (n = 136), respectively (Fig. 2b). Beyond matching the trajectories of subjects (Fig. 2c, Extended Data Fig. 1; Task 1: β = 2.58, p = 3.024 × 10^−40^; Task 2: β = −0.53, p = 6.00 × 10⁻⁴; linear mixed-effects model), the Full model, evaluated under this generalization condition without additional training, achieved a 55.14% success rate (n = 1478) (Fig. 2d), exceeding subject-level performance (χ²(1) = 52.53; p = 4.231 × 10⁻¹³; chi-squared test). In contrast, the No-RS model achieved only a 5.59% success rate (n = 1485), performing substantially worse than both the structured model and macaque subjects (χ²(1) = 409.56; p = 4.563 × 10^−91^; chi-squared test). To assess whether the Full model’s performance depended on the correspondence between entities and their edge weights, we exchanged the prey and predator edge weights during evaluation without rearranging node features or modifying learned parameters. This manipulation reduced success from 55.14% to 22.18% (Fig. 2d; t(29) =-18.764; p = 9.171 × 10^−18^; two-tailed paired t-test), indicating that the original edge-weight correspondence contributes to performance. Together with the Full and No-RS model comparison, this finding suggests that the absence of explicit graph-based aggregation in No-RS may contribute to its poorer zero-shot generalization. Consistent with this, the model also generalized to changes in environmental dynamics (e.g., arena size or friction) and prey properties (e.g., speed or number) (Extended Data Fig. 1).

The relational structure model predicts that entities are represented in a common relational format. Each entity is processed through the same set of transformations according to its relation to the agent, rather than its fixed identity. We hypothesized that dACC might show a comparable property. To test this, we trained a decoder on the relative distance between the subject and one entity, and evaluated its ability to generalize to a different entity. The decoder was able to generalize in both directions between the two preys (prey 1 → prey 2: 37.8%, z = 11.6, p = 2.7 × 10⁻³¹; prey 2 → prey 1: 39.2% at 50–87 pixels, z = 13.6, p = 2.8 × 10^−42^; right tailed z-test; chance = 10 %; Fig. 2e), indicating that relative distance, for preys, are encoded in a format that generalizes across entities. Cross-decoding also generalized between predator and prey (prey → predator: 58.5%, z = 26.5, p = 1.2 × 10^−154^; predator → prey: 36.0%, z = 15.3, p = 3.5 × 10^−53^; Fig. 2e), suggesting that this representation is not specific to prey and extends to an entity with a different behavioral role. Although the trend showed that cross-decoding was more significant at short distances near capture, decoding was significant across multiple distance bins. In contrast, the same decoder could not decode the distance to the non-pursued prey with the same procedure (prey 1 → prey 2: 19.8% vs 20.2% surrogate mean, z = −0.21, p = 0.58; prey 2 → prey 1: 32.0% vs 33.5%, z = −0.82, p = 0.80; Fig. 2f). Within trials, the distance between gaze and the pursued prey decreased as the self approached it. The within-trial correlation between the relative distance and gaze-prey distance was positive in 86% of the trials (median Spearman ρ = 0.26; Wilcoxon signed-rank test, z = 36.0, p = 6.8 × 10^−284^, n = 2,304 trials; Extended Data Fig. 5b). Thus, gaze behavior was increasingly aligned with the pursued prey as the subject approached it, providing behavioral context for the selectivity of neural cross-decoding. Together, these findings identify a corresponding relational structure in dACC, consistent with the model’s prediction that relative spatial relationships are represented in a common format across entities with different identities and behavioral roles.

### Selective attention supports stable target selection as the number of prey increases

In a multi-entity pursuit scenario, agents encounter entities with distinct behavioral policies and action possibilities. As the number of entities increases, encoding all entities simultaneously leads to a combinatorial explosion of information. We therefore hypothesized that selective attention (i.e., spotlight attention) mitigates this problem and constrains effective dimensionality. To examine the contribution of attentional selectivity, we increased the edge-entropy coefficient in the comparison models (Fig. 3a). Increasing this coefficient encourages more uniform attention weights, thereby weakening selective attention.

**Fig. 3.**
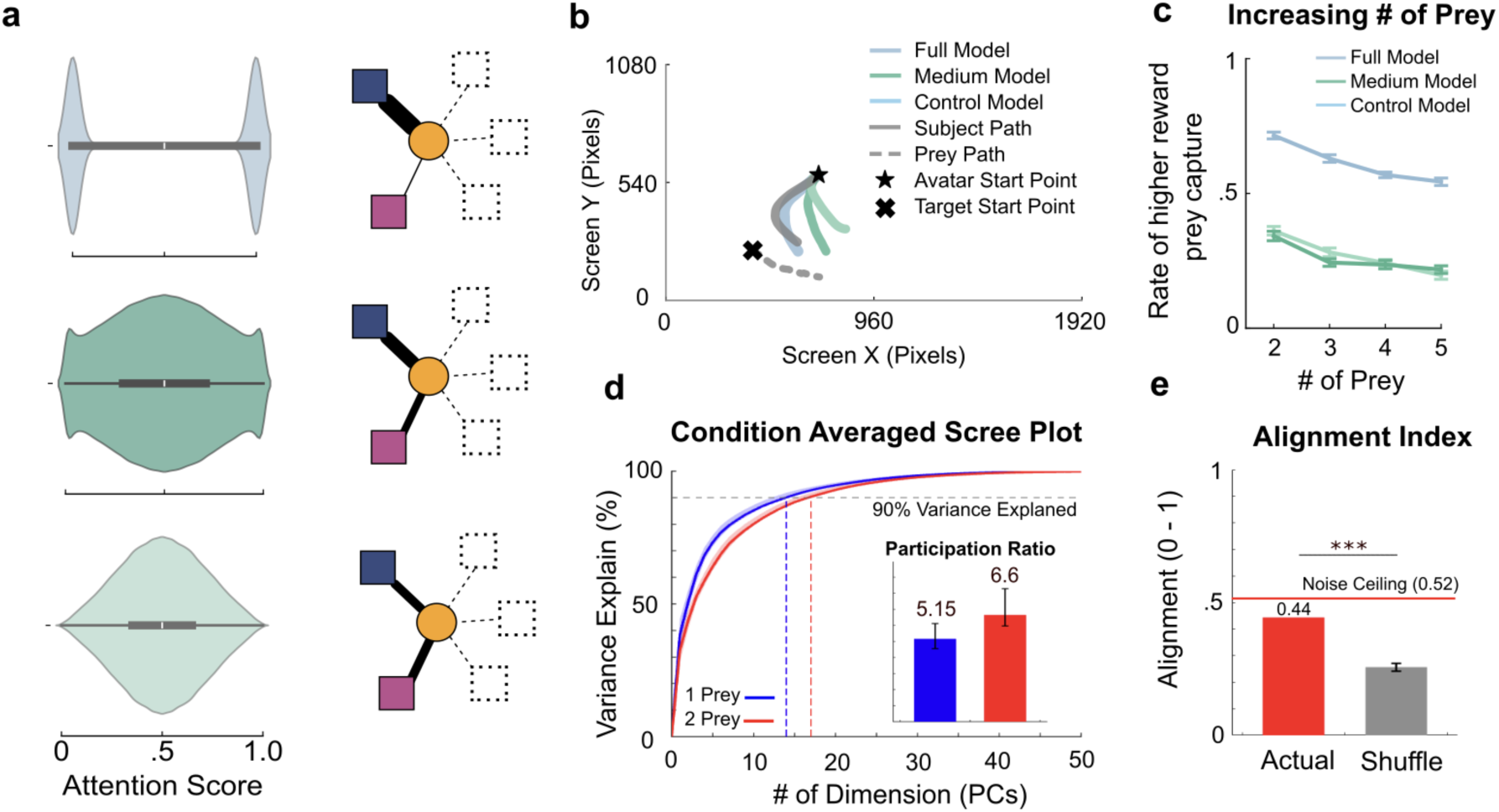
Spotlight Attention Results and the Neural Implementation of Dimensionality. **a)** Attention-weight distribution and corresponding weighted graphs for the low (Full model; sky blue), medium (Medium-EC model; green), and high (High-EC model; light green) edge-entropy coefficient conditions. In the graph diagrams, edge thickness indicates the weight assigned to each target. Higher coefficients encourage more uniform weights, attenuating selective attention. b) Example trajectories and corresponding model-generated trajectories from different ablation models, compared with subject behavior. c) Success rate for intercepting higher-reward prey as a function of the number of targets. The Full model is shown in skyblue, with the Medium-EC and High-EC models in green and light green, respectively. d) Scree plot of explained variance from condition-averaged neural activity during the first second of the single-prey (blue) and double-prey (red) conditions. The x-axis denotes the number of principal components (PCs), and the y-axis denotes explained variance. Dashed lines indicate the number of PCs required to explain 90% of the variance. Inset: participation ratio for each condition, reflecting effective dimensionality. Error bars denote s.e.m. e) Alignment index between neural subspaces. The top 10 PCs from the double-prey condition are projected onto those from the single-prey condition (red), compared with shuffled data (grey). The red line indicates the noise ceiling from split-half reliability. (* p < .05, ** p < .01, *** p < .001)

As expected, the model with spotlight attention mimicked subjects’ behavior better than models with distributed attention (Fig. 3b; Extended Data Fig. 2; Medium edge-entropy coefficient(Medium-EC) - Task 1: β = 1.25; p = 9.247 × 10^−13^; task2: β = 0.70; p = 2.575 × 10^−5^; High edge-entropy coefficient (High-EC) - Task 1: β = 1.47; p = 5.947 × 10^−17^; task2: β = 0.90; p = 5.211 × 10^−8^; linear mixed-effects model). To further investigate the necessity of spotlight attention in flexible control, we increased the number of prey the agent encountered in each trial. Increasing the number of options also increases the demand to encode multiple entities with distinct actions and behavioral policies. The model with spotlight attention (i.e., low regularization) consistently outperformed the other models as the number of prey increased (Fig. 3c; vs High-EC model (high regularization; uniform attention), 2: t(55.16) = 17.51; p = 3.440 × 10^−24^; 3: t(56.79) = 16.34; p = 3.964 × 10^−23^; 4: t(54.59) = 19.38; p = 3.837 × 10^−26^; 5 prey: t(57.65) = 17.09; p = 3.105 × 10^−24^; Welch’s t-test). Only through selective attention can we observe that the agent maintains a stable commitment to pursuing higher-reward prey across all other generalization conditions (Extended Data Fig. 2). These results demonstrate that spotlight attention effectively addresses the combinatorial explosion by constraining the encoding space, thereby enabling stable commitment as environmental complexity increases.

If similar computational principles are implemented in biological circuits, neural population dimensionality should remain stable despite increases in the number of competing entities. Indeed, we found that population dimensionality was comparable across the two tasks (Fig. 3d), with similar participation ratios despite the increase in available options (Fig. 3d inset; task 1 = 5.15; task 2 = 6.60). Notably, comparable dimensionality leaves open the possibility that different representational axes or dynamics are engaged across conditions, which is known as neural reuse. To test this, we compared the neural subspaces across tasks. The subspaces were strongly aligned (Fig. 3e; z = 12.71; p = 2.460 × 10^−37^; right-tailed z-test), indicating that the dACC population reuses a common representational geometry rather than recruiting distinct prey-specific subspaces. This alignment suggests that, within dACC, activity during two-prey pursuit remains within the same representational basis used for single-prey pursuit. Together, these findings support the idea that spotlight attention enables scalable, flexible control by compressing information into a shared low-dimensional subspace that generalizes across contexts.

### Computing affordance resolves encountering a prey that exceeds the agent’s own physicality

Pursuit is an embodied behavior in which success depends on the relative physical relationships between the agent and its targets, including distance, speed, and heading direction^38^. Consequently, targets with identical rewards can differ in their actionability depending on their physical configuration. Hence, models that fail to represent these relationships are expected to generalize poorly under changing conditions. We hypothesized that flexible control depends on encoding such relative structure, enabling the agent to evaluate feasibility and redirect pursuit when necessary.

To test this, we compared a model that learned to compute relative physical information, namely affordance, with one that relies only on reward-based features (Fig. 4a). As expected, the model with learned affordance scores (affordance-based model) more closely matched subjects’ behavior than the reward-only model (Fig. 4b; Extended Data Fig. 3a; Task 1: β = 0.39, p = 1.87 × 10⁻²; Task 2: β = 0.51, p = 2.30 × 10⁻³; linear mixed-effects model), despite the latter achieving comparable or higher reward rates under standard conditions (Extended Data Fig. 3, Extended Data Fig. 4). Thus, maximizing reward did not by itself produce subject-like behavior. Consistent with the view that alignment between artificial and biological systems depends on shared constraints rather than on a shared objective alone^39^, the two models diverged most clearly in conditions where physical feasibility constrained behavior. A potential concern was that similar behavior could arise from different underlying representations. Therefore, we computed TTC-affordance for the model and the macaque to post-hoc compare the two affordance maps. The affordance map representations were closely aligned between the model and subjects (Fig. 4c; mean cosine similarity = 0.64; 95% CI = [0.63, 0.65]; p = 5.00 × 10^−6^; sign-flip permutation test). Notably, under standard conditions, both models could successfully capture prey, suggesting that reward-based strategies may suffice when physical feasibility is not limiting. To directly test the necessity of the learned affordance scores, we introduced a valuable but uncatchable prey whose speed exceeded that of the agent. Under this condition, the No-affordance model exhibited near-zero success compared to the Full model (Fig. 4d; t(33.76) = 34.01; p = 1.043 × 10^−27^; Welch’s t-test), indicating a failure to account for physical constraints. Together, these results demonstrate that affordance-based commitment is more robust than reward-based selection when physical feasibility is critical, enabling flexible disengagement and redirection of pursuit.

**Fig. 4.**
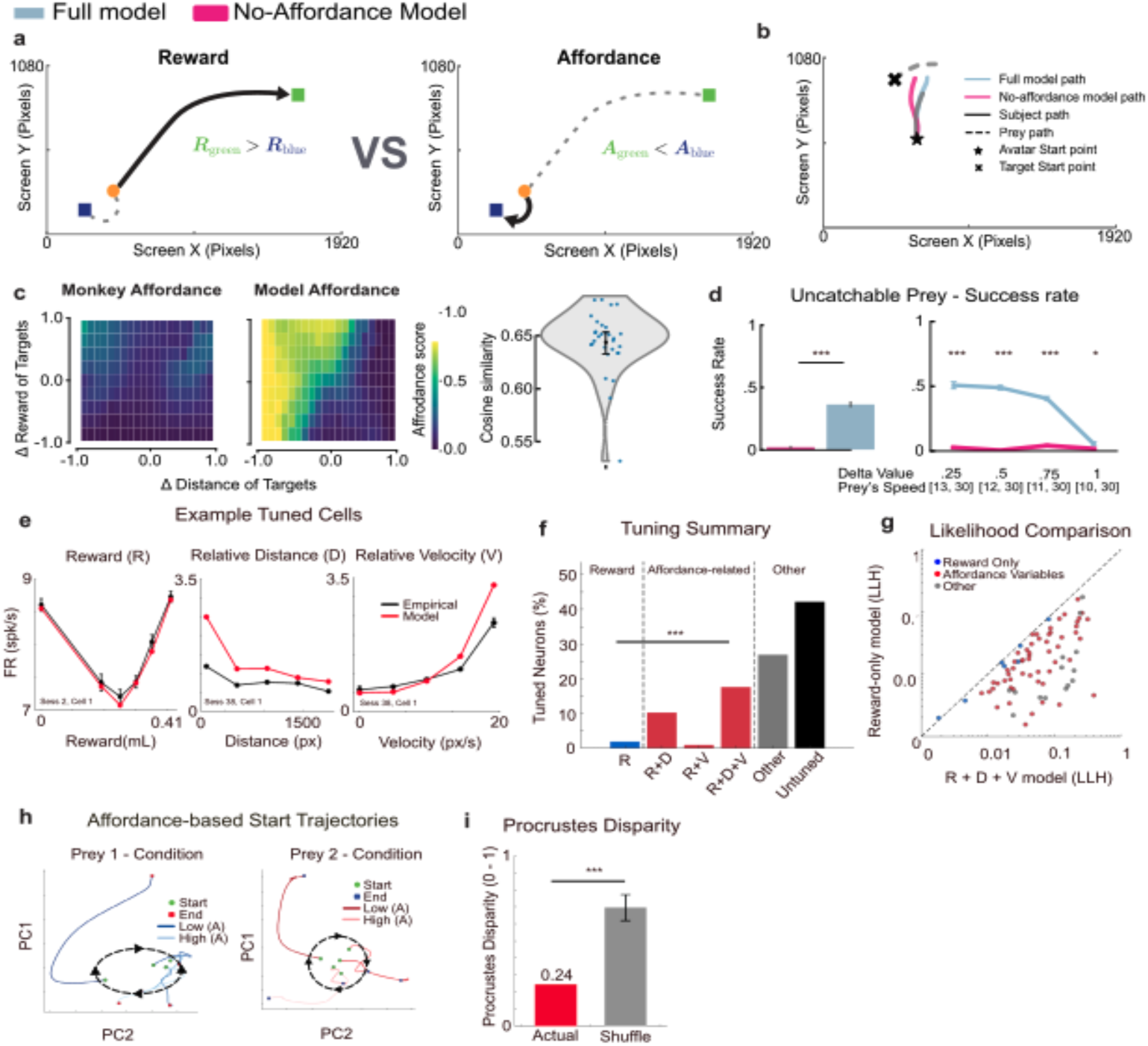
Affordance Computation Results and the Neural Implementation of Affordance Tuning. **a)** Comparison of the pursuit paths of the reward-based No-affordance model and the affordance-based Full model. R denotes reward value, and A denotes learned affordance scores. b) Example corresponding fit trajectories of the Full model and the No-affordance model. c) TTC-Affordance map of the subject (left) and model (center). The z–axis represents the affordance score. (right) Cosine similarity between the subject’s and model’s affordance structure. d) Success rate of the No-affordance and Full model in the uncatchable-prey condition, shown overall (left) and across parameter settings (right). e) Single neuron example of a tuned neuron to variable reward denoted as R, relative distance denoted as D, and relative velocity denoted as V. The red line represents the model prediction while the black line shows the empirical firing rate with SEM±1 bar at each bin. f) Percentage of neurons significantly tuned (q < 0.05, corrected for multiple comparisons). Categories include reward-only tuning shown in blue, affordance-related variables shown in red (Reward + Distance (RD), Reward + Velocity (RV), Reward + Distance + Velocity (RDV) model), other variables tuning shown in grey (Velocity-only (V), Distance-only (D), and Velocity + Distance (VD) model), the other category also includes untuned cells in black. g) Likelihood value comparison where each dot represents significantly tuned cells, and they are color-coded the same as panel F. The y-axis represents the LLH value for the reward-only tuned model, while the x-axis represents RDV LLH values. h) Principal Component Analysis (PCA) trajectory plots of the two different conditions of the first 0.5 seconds of each trial. The light blue and light red indicate high TTC-affordance (aff > 0.5269, aff > 0.6603, respectively), while the dark blue and dark red indicate low affordance (aff < 0.1316, aff < 0.1554, respectively). The black dashed circle indicates an orderly progression of affordance values around the circle, from low to high. i) Procrustes disparity between neural population dynamics in Task 1 and Task 2. Red indicates observed neural data; grey indicates shuffled data. Error bars are the s.e.m. (* p < .05, ** p < .01, *** p < .001)

To test whether dACC encodes affordance, we examined neuronal tuning to both its constituent variables (e.g., distance and angle to prey) and their composite. We constructed a composite TTC-based affordance variable that integrates relative physical information, reward magnitude, and moment-to-moment reward discounting (see Methods), and estimated neuronal selectivity using a linear–nonlinear Poisson (LNP) model (Fig. 4e). We found that about 57.84% of the neurons were tuned to one or more variables (q < 0.05; Benjamini-Hochberg FDR-adjusted p-value), only 2% being tuned to reward-only (R) compared to 17.65% being tuned to reward, distance, and velocity (RDV) (Fig. 4f; p = 9.96 × 10^−8^; two-sided binomial test). We additionally examined the log-likelihood (LLH) values of the R and RDV models. The graph showed a skewness towards the RDV model, implying that individual neurons are more tuned to affordance-related variables than to reward itself (Fig. 4g). We next examined whether the population encodes TTC-affordance values by projecting activity into a low-dimensional subspace using principal component analysis (PCA). When population activity from 1-prey and 2-prey conditions was projected into this space, trajectories corresponding to different affordance states occupied distinct regions of the subspace (Fig. 4h). To determine whether the TTC-affordance-dependent dynamics remain invariant between 1-prey and 2-prey conditions, we quantified trajectory similarity between the conditions using Procrustes analysis. The distance between empirical dynamics was significantly smaller than that obtained from shuffled data (Fig. 4i; d = 0.25; z = −5.89; p = 1.910 × 10^−9^; left-tailed z-test), indicating that the affordance shaped the shared geometric structure across conditions. Together, these findings suggest that dACC population activity reflects multiple task-relevant factors, including reward and physical properties of potential targets, and that these variables are represented within a shared population geometry during pursuit. Although whether the macaque brain implements the specific affordance computation proposed by our model remains an open question, these findings motivate the conceptual hypothesis that reward and physical feasibility are jointly represented and may contribute to evaluating pursuit opportunities.

### Change of Mind as Emergent Testbed and Neural Substrates in Biological Agents

As an emergent behavior in biological agents, change-of-mind (CoM) reflects the dynamic re-evaluation of competing options and provides a natural test of real-time adaptive control^40^. Because CoM arises across a wide range of relational and affordance configurations, it provides a mechanistic probe of flexible generalization beyond specific task settings. Compared to the preceding analyses, CoM constitutes a more stringent test, as it unfolds in real time and is not directly optimized during training. We therefore hypothesized that agents equipped with all three cognitive constructs would reproduce CoM behavior comparable to that of biological agents. In our paradigm, we defined CoM trials as instances in which the agent redirected pursuit from one prey to another as action feasibility evolved (Fig. 5a).

**Fig. 5.**
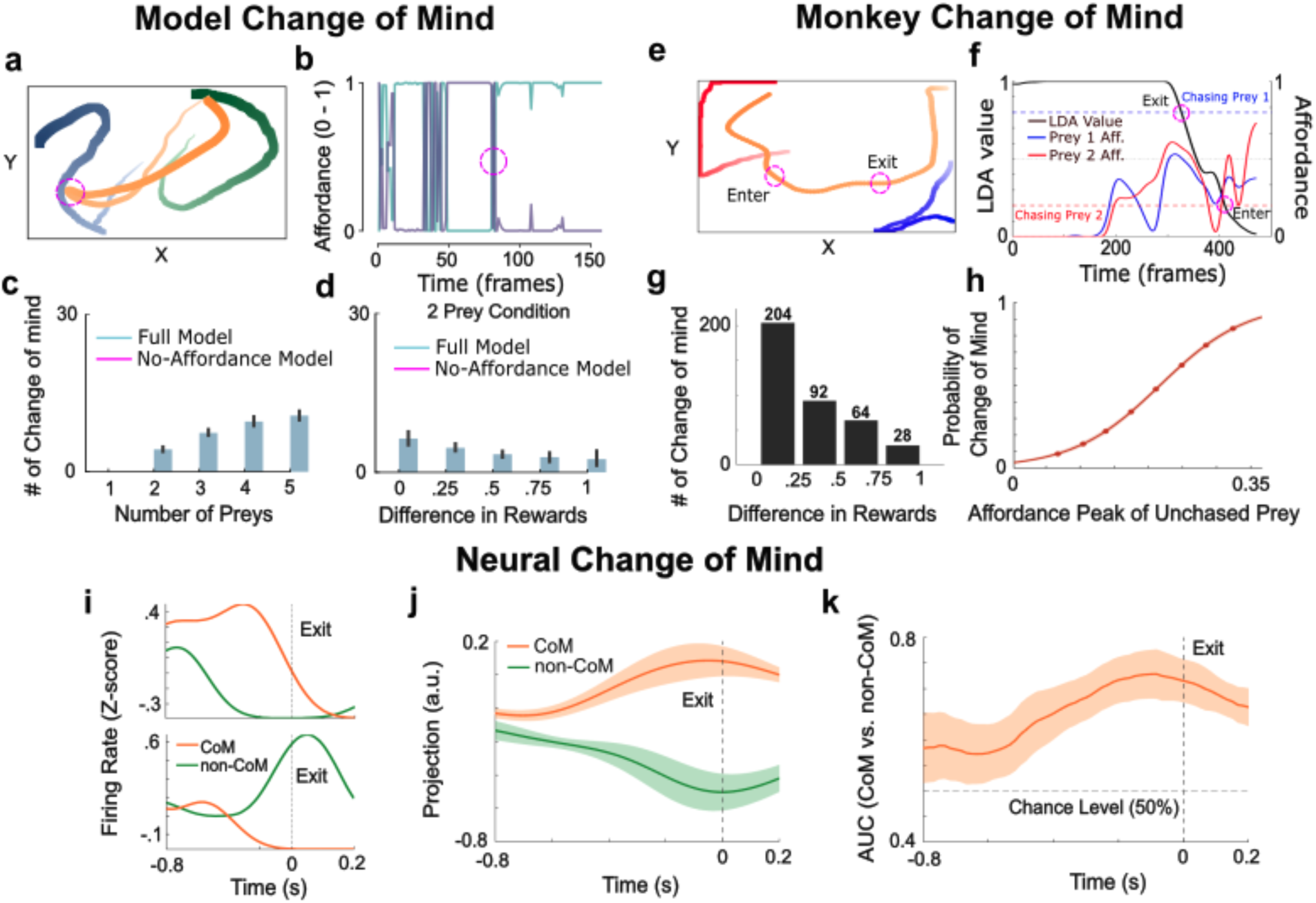
Comparison of the Number of ‘Change of Mind’ Events Between the Full model and the No-affordance model Across Different Numbers of Prey. **a)** Example artificial agent CoM trial. The agent is shown in orange; prey are blue and green. The avatars become progressively larger over time for visual purposes. b) Example affordance traces from the agent; the magenta marker indicates the moment of CoM. c) CoM counts for Full (skyblue) versus No-affordance (magenta) models across increasing prey numbers. d) CoM counts based on reward difference in the 2-prey condition. e) Example subject CoM trajectory. f) Linear discriminant analysis (LDA) model predicted state and affordance traces for a representative trial. The black line is the LDA decision variable; dashed blue/red lines indicate LDA confidence bounds for commitment; solid blue/red lines are affordances for prey 1 and prey 2. g) CoM trial counts versus normalized reward difference (CoM trials, n = 388). h) Psychometric curve of the average of both subjects. The probability of Change of Mind is a function of the peak affordance of the non-initially chased prey before the switch. i) Single-neuron examples (z-scored, aligned to exit). The orange line denotes CoM trials, and the green line denotes non-CoM trials. j) Projection of population activity onto the CoM coding axis (difference of CoM vs non-CoM means at exit), plotted over time. k) Cross-validated AUC over time from the CoM coding axis.

Consistent with this prediction, the model exhibited CoM behavior despite not being explicitly trained to do so. During CoM trials, the artificial agent was able to reallocate attention from the initially selected prey to an alternative target and subsequently commit to the new pursuit (Fig. 5b). Therefore, to examine the conditions under which CoM behavior emerges, we manipulated factors expected to increase fluctuations. Increasing the number of prey in the scene introduces greater competition and more frequent shifts in the relative feasibility of actions. Consistent with this prediction, the model exhibited a higher rate of mid-course reversals as prey number increased (Fig. 5c; β = 2.68 ± 0.12; p = 1.783 × 10^−48^; linear regression). In contrast, the reward-only model, which cannot generate learned affordance scores, rarely reversed its decisions. Similarly, when the reward gap between prey was small, resulting in more frequent shifts in relative affordance, the model exhibited increased reversal frequency (Fig. 5d; β = −3.83 ± 0.68; p = 8.957 × 10^−8^; linear regression). Together, these results indicate that CoM behavior scales with affordance instability rather than reward magnitude alone.

To quantify change of mind (CoM) events in subjects, we trained a linear discriminant analysis (LDA) classifier with fifty moment-to-moment kinematic features of all entities present within a trial (Fig. 5e; Extended Data Fig. 6; see Methods). Then we used the LDA scores from the logistic readout to compute the posterior probability that the subject was pursuing either prey at every frame. We defined the subject as committed to a prey whenever this posterior lay within 15% of the corresponding bound (p < 0.15 for prey 1, p > 0.85 for prey 2). We classified a trial as a CoM trial when the subject committed to the prey it did not ultimately capture for at least 0.5 s (30 consecutive frames), and this commitment ended before the first frame of commitment to the captured prey. We defined the reversal onset as the exit, the first frame at which the classifier started drifting away from the initially chased prey. In contrast, we defined reversal completion (entrance) as the first frame of commitment to the new prey. In parallel, we computed a moment-to-moment TTC-affordance value for each prey, yielding a continuous measure of action feasibility. This allowed us to test whether CoM was driven by the TTC-affordance or other variables (Fig. 5f; see Methods). Consistent with the model, subjects reversed more often when the reward gap between prey narrowed (Fig. 5g; β = −3.04 ± 0.48; p = 2.2 × 10⁻¹⁰; logistic regression). Together, these findings indicate that macaque CoM behavior is related to the changing feasibility of alternative targets, supporting a prediction of the affordance-based model, while the precise computational mechanisms underlying this relationship in biological agents remain open.

We then directly tested whether TTC-affordance dynamics predicted reversals by fitting a weighted logistic regression model using the peak affordance of the non-pursued prey prior to the reversal. The probability of reversal increased with this TTC-affordance peak (Fig. 5h; p = 7.49 × 10^−205^; weighted logistic regression; see Extended Data Fig. 7 for individual subject results). Together, these findings indicate that CoM behavior in biological agents is related to affordance signals, consistent with the prediction of the model.

We next asked whether the dACC population activity reflects the re-evaluation process underlying CoM. At the single-neuron level, responses around reversals were heterogeneous and mixed (Fig. 5i; Extended Data Fig. 8), suggesting that CoM is better understood at the population level. We therefore defined a CoM coding axis in the population as the normalized difference in mean activity between CoM and non-CoM trials at exit time (Fig. 5j). Projecting exit-aligned activity onto this axis revealed a clear separation between CoM and non-CoM trials near the moment of disengagement from the initially pursued prey. To assess generalizability, we performed leave-one-session-out cross-validation and found significant decoding of CoM over - 0.8 seconds before and 0.2 seconds after switch onset, with peak decodability around exit (Fig. 5k; AUC = 0.73; p = 0.0312; one-sided Wilcoxon signed-rank test). These results indicate that dACC population activity contains reliable information about impending behavioral switches, consistent with a population-level signal tracking the reallocation of pursuit as the affordance landscape evolves.

## Discussion

Previous studies on zero-shot generalization have often emphasized either the training curriculum^41^, representational constraints^42^, or structural priors that constrain the hypothesis space^20^. We extend this perspective by showing that several classic cognitive constructs—relational structure, spotlight attention, and affordance computation—jointly enable generalization in embodied, closed-loop control, where feasibility evolves moment-to-moment rather than remaining fixed, unlike traditional cognitive neuroscience tasks. Within this architecture, removing a construct impaired a different form of generalization: robust transfer and biologically characteristic decision dynamics emerged only when these mechanisms operated in concert. Such dissociation suggests that, within this model, the three constructs contribute differently to zero-shot transfer. More broadly, this work illustrates how integrating multiple cognitive constructs within a single architecture enables the study of complex behavior, moving beyond the common practice of modeling isolated functions with neural networks^43–46^.

An important distinction concerns how we interpret the reward-only model and the role of learning in our task. Through repeated interactions and sufficient training, a sufficiently expressive reinforcement-learning agent, including a deep RL system such as DQN, could potentially learn to avoid pursuing physically infeasible targets through reward feedback alone. Our results do not suggest an inherent limitation of reward-based reinforcement learning. Rather, our comparison addresses a different question: whether an agent can generalize its target selection to novel physical conditions without additional training. In this zero-shot setting, physical conditions change while the relative reward values of the targets remain unchanged. Our No-affordance model determines target selection and constructs graph connections based on reward magnitude, without explicitly incorporating position and velocity into the evaluation of physical pursuability. Consequently, this component lacks an explicit mechanism to adapt target selection to changing physical feasibility. In contrast, the Full model incorporates position, velocity, and reward into its affordance computation, allowing target evaluation to adapt dynamically to physical conditions. Thus, our findings highlight the contribution of explicitly incorporating physical actionability to zero-shot generalization in the tested task, rather than establishing a general limitation of reward-based reinforcement learning.

Several limitations still constrain our conclusions. First, our model assumes full access to the environment state, whereas real-world pursuit is typically partially observable and better described as a partially observable Markov decision process (POMDP), requiring belief maintenance and active information sampling rather than direct access to the state^47,48^. Biological agents reduce uncertainty through selective sensing (e.g., gaze shifts and foveated vision), processes that our current framework does not implement^49^. Extending the model to include egocentric, limited observations with explicit belief-state inference and active sensing would provide a stronger test of whether the same components remain sufficient under partial observability. Second, although the affordance scores are learned from reward rather than the specified analytical affordance, we chose the variables, position, velocity, and reward, available to the affordances module in advance. This introduces prior knowledge about the task and constrains the types of affordances the model can represent. Accordingly, the model does not establish how biological agents acquire such computations, nor whether they use the specific calculation proposed here. Rather, our results motivate the conceptual hypothesis that reward and physical feasibility may be jointly represented when evaluating pursuit opportunities. Learning to discover affordances from experience, including affordances relevant in different environments, is an important direction for future work. Third, our neural analysis focuses exclusively on the dorsal anterior cingulate cortex (dACC), offering only a partial view of the circuitry underlying attention, relational structure, and affordance-like prioritization. These computations are likely distributed across sensory, attentional, sensorimotor, valuation, and action-selection networks. Future work with multi-region recordings will clarify how these representations interact and whether dACC acts as an integration hub or reflects computations formed elsewhere.

Traditional reverse engineering has followed two main paths: fitting behavioral data with latent process models and searching for neural correlates of inferred parameters^50,51^, or reproducing observed neural dynamics by imposing architectural constraints or modifying learning rules in network models to match them^44,52^. Both strategies assume that the relevant computational motifs can be parameterized a priori and that each motif independently shapes behavior. However, in complex, closed-loop tasks such as dynamic pursuit, it is difficult to derive a simplified latent process or isolate a single defining computational motif that fully captures the underlying neural dynamics. In this regard, our use of network models suggests a complementary approach to reverse-engineering such complex behavior. We use artificial agents as computational testbeds to evaluate multiple candidate architectural constraints and representations when the underlying decomposition is uncertain. More recently, related approaches have embedded modular learning structures to adjudicate among competing memory hypotheses^53^. Here, we extend this strategy by grounding separable computational motifs in classical cognitive theory, integrating them into a unified architecture, and testing their necessity for zero-shot transfer and process-level behaviors not explicitly optimized during training, such as change-of-mind dynamics. Importantly, the correspondence between model-derived predictions and biological behavior provides a means of evaluating candidate computational accounts, rather than establishing that biological agents implement the model’s specific architecture. In the case of CoM, the affordance-based mechanism predicts that pursuit reversals should be related to changes in the relative feasibility of competing targets. The macaque behavioral analysis provides evidence consistent with this prediction, showing that reversal probability is associated with the TTC-based analytical affordance of the alternative prey. This correspondence suggests that changes in target feasibility may contribute to flexible pursuit behavior, although the precise computational mechanisms underlying this relationship in biological agents remain to be determined. Similarly, our neural analyses identify population-level properties that can be compared with representational features of the model, providing a basis for further investigation of their potential relationship. More broadly, this framework illustrates how computational models can help formulate and examine hypotheses about complex biological behavior, extending the study of computational principles beyond tightly controlled paradigms.

## Extended Figures

**Extended Data Fig. 1.**
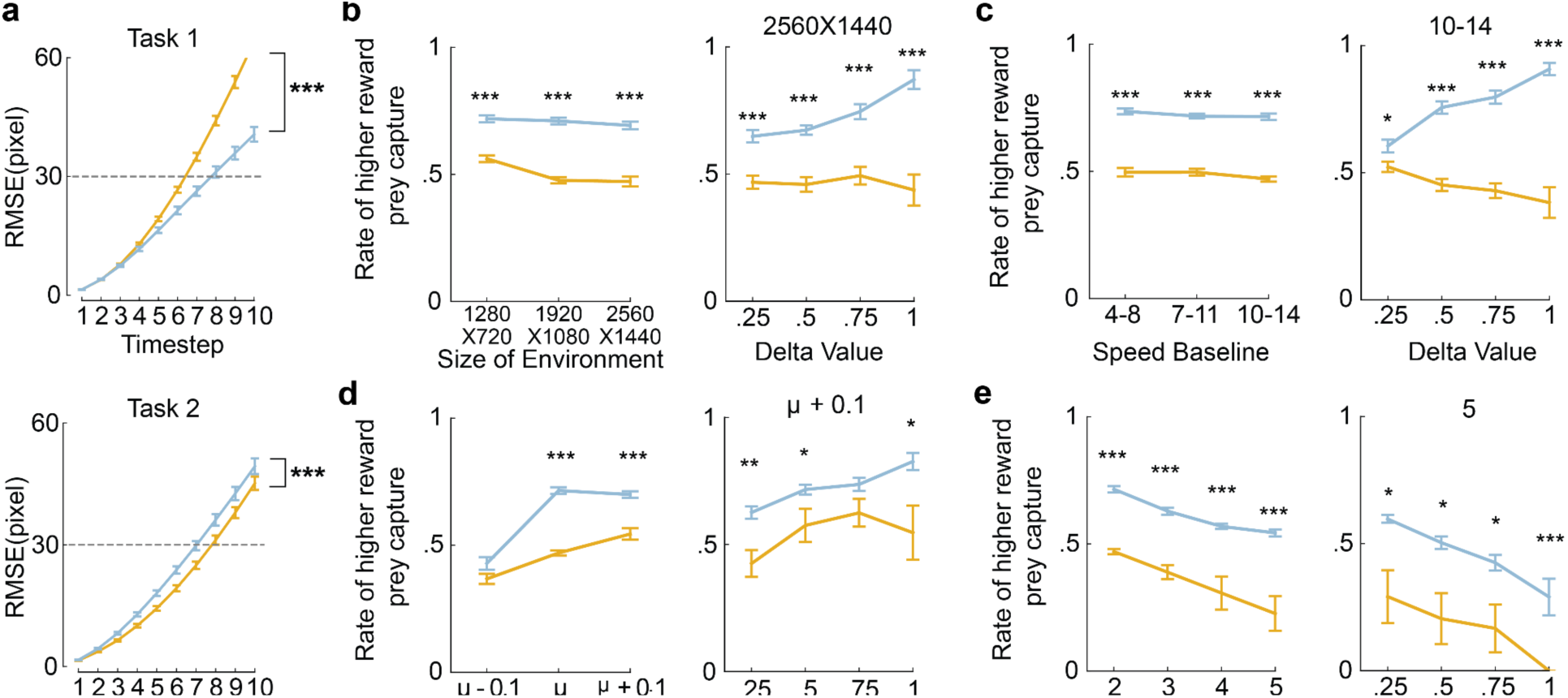
Ablation of Relational Structure Results. **a)** Comparative RMSE between models of the rollout trajectory prediction in Task 1 (left) and 2 (right). b) Rate of higher-reward prey capture across different environment sizes (left) and as a function of the target value difference in the biggest size (right). c) Rate of higher reward prey capture across different target speeds (left) and as a function of the target value difference at the highest speed (right). d) Rate of higher reward prey capture across different friction levels (left) and as a function of target value difference in the increased friction (right). e) Rate of higher reward prey capture across different numbers of targets (left) and as a function of target value difference in the five prey (right).

**Extended Data Fig. 2.**
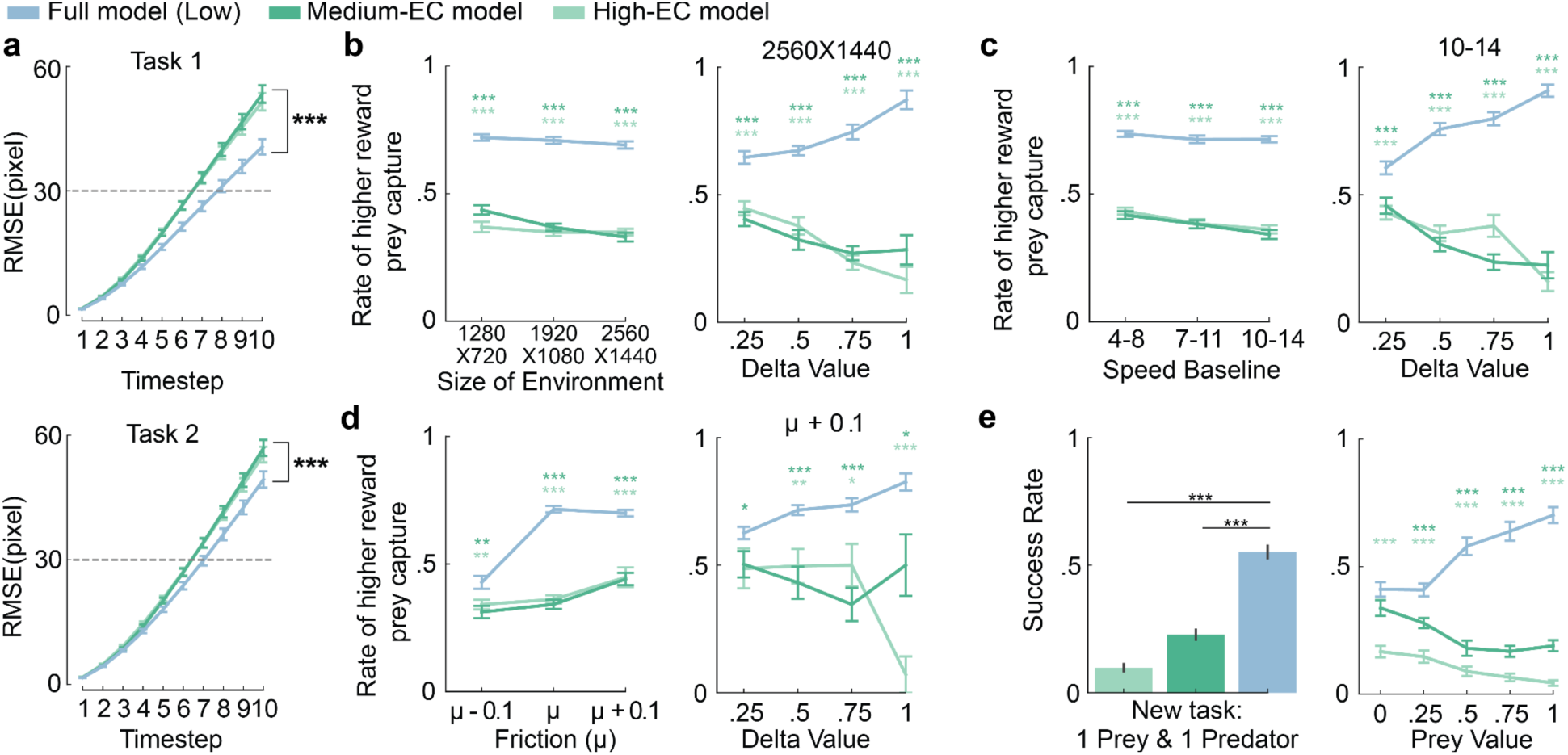
Ablation of Spotlight Attention Results. **a)** Comparative RMSE between models of rollout trajectory prediction in Task 1 (top) and 2 (bottom). b) Rate of higher reward prey capture across different environment sizes (left) and as a function of the target value difference in the biggest size (right). c) Rate of higher reward prey capture across different target speeds (left) and as a function of the target value difference at the highest speed (right). d) Rate of higher reward prey capture across different friction levels (left) and as a function of target value difference in the increased friction (right). e) Success rate in the ‘1 prey, 1 predator’ scenario (left) and as a function of target value difference (right).

**Extended Data Fig. 3.**
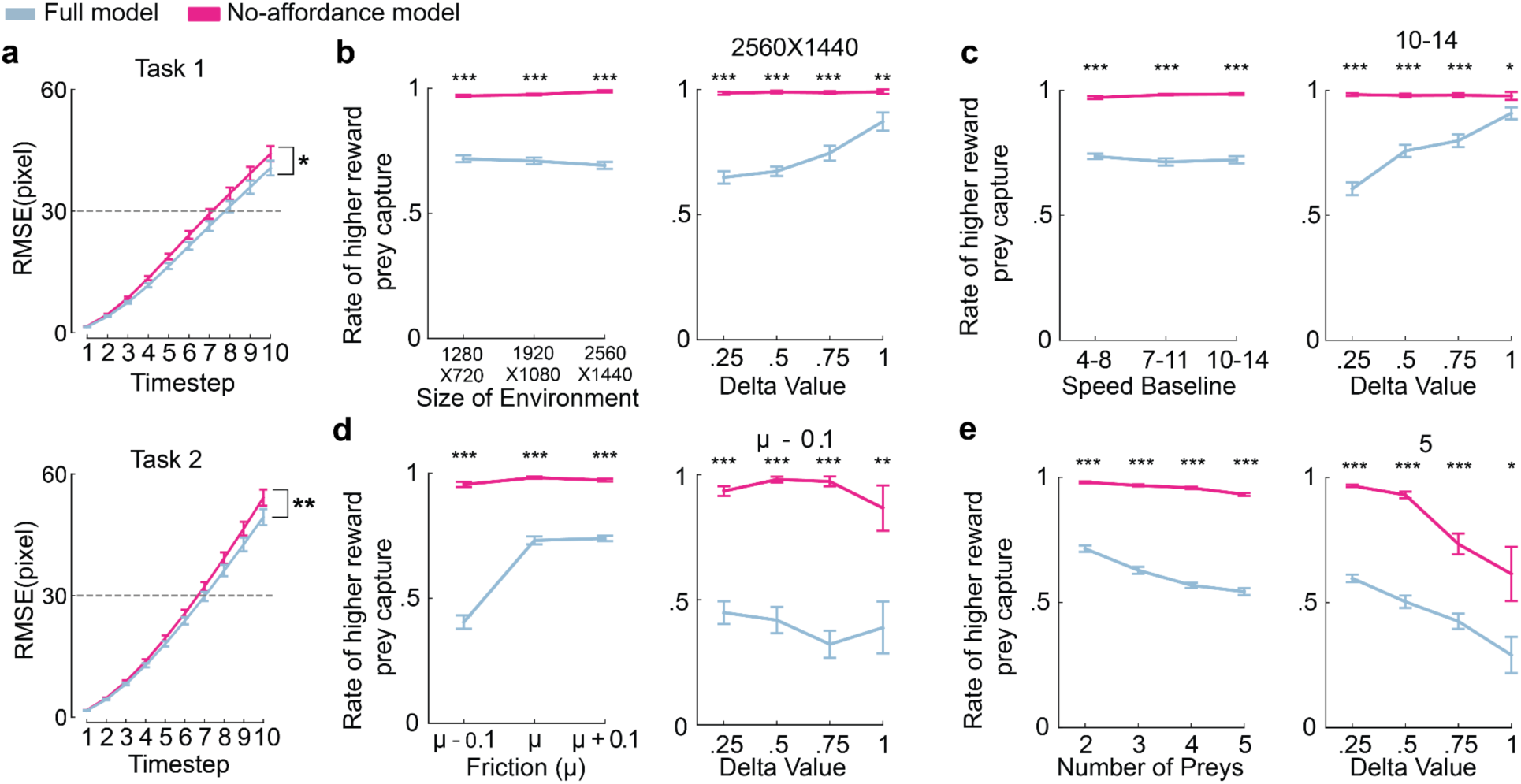
Results of Affordance Ablation. **a)** Comparative RMSE between models in rollout trajectory prediction in Task 1 (top) and 2 (bottom). b) Rate of higher reward prey capture across different environment sizes (left) and as a function of the target value difference in the biggest size (right). c) Rate of higher reward prey capture across different target speeds (left) and as a function of the target value difference at the highest speed (right). d) Rate of higher reward prey capture across different friction levels (left) and as a function of target value difference in the decreased friction (right). e) Rate of higher reward prey capture across different numbers of targets (left) and as a function of target value difference in the five prey (right).

**Extended Data Fig. 4.**
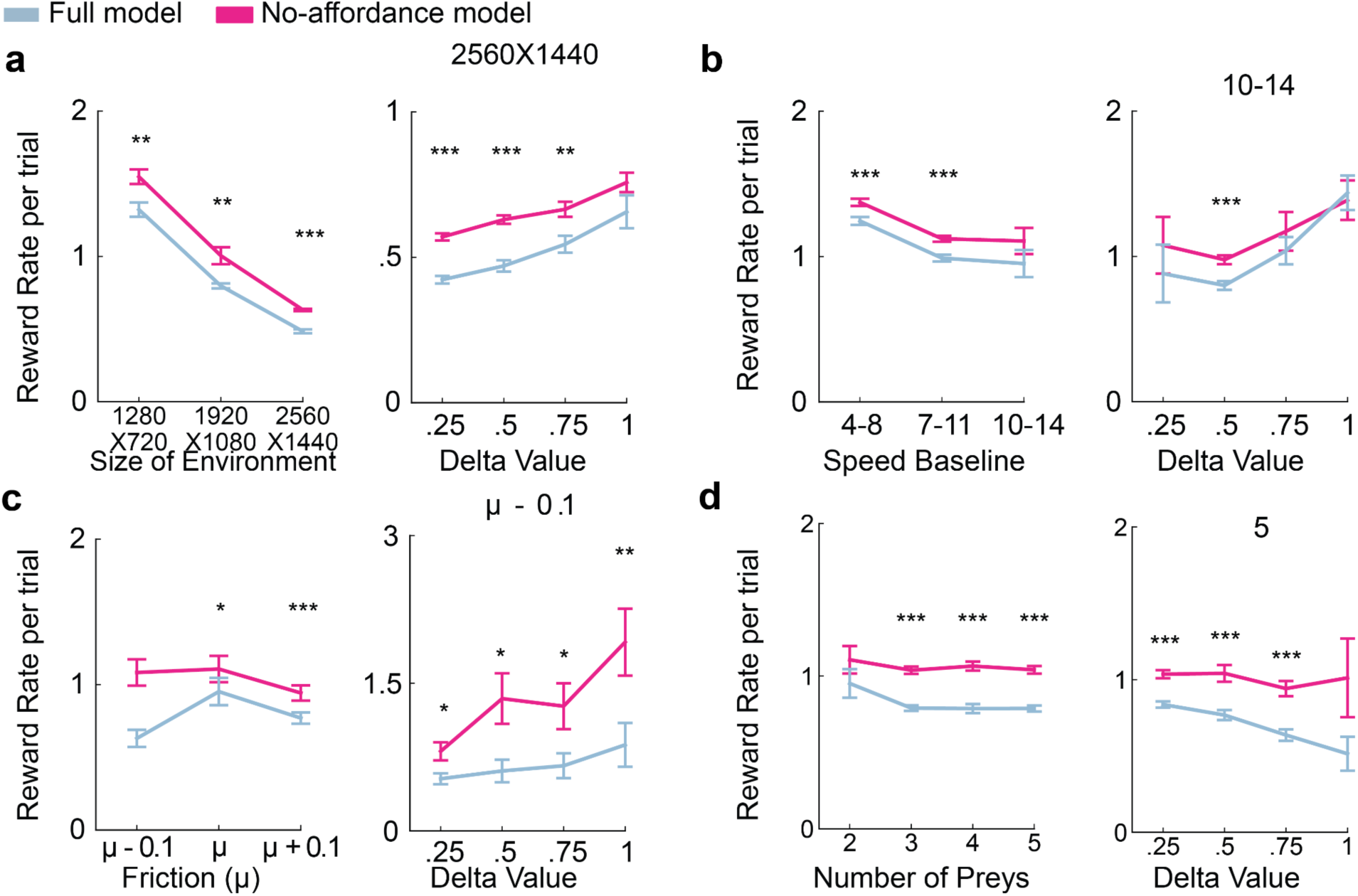
Affordance Model Reward Rate Results. **a)** Reward rate per trial across different environment sizes (left) and as a function of the target value difference in the biggest size (right). b) Reward rate per trial across different target speeds (left) and as a function of the target value difference in the highest speed (right). c) Reward rate per trial across different friction levels (left) and as a function of target value difference in the decreased friction (right). d) Reward rate per trial across different numbers of targets (left) and as a function of target value difference in the five prey (right).

**Extended Data Fig. 5.**
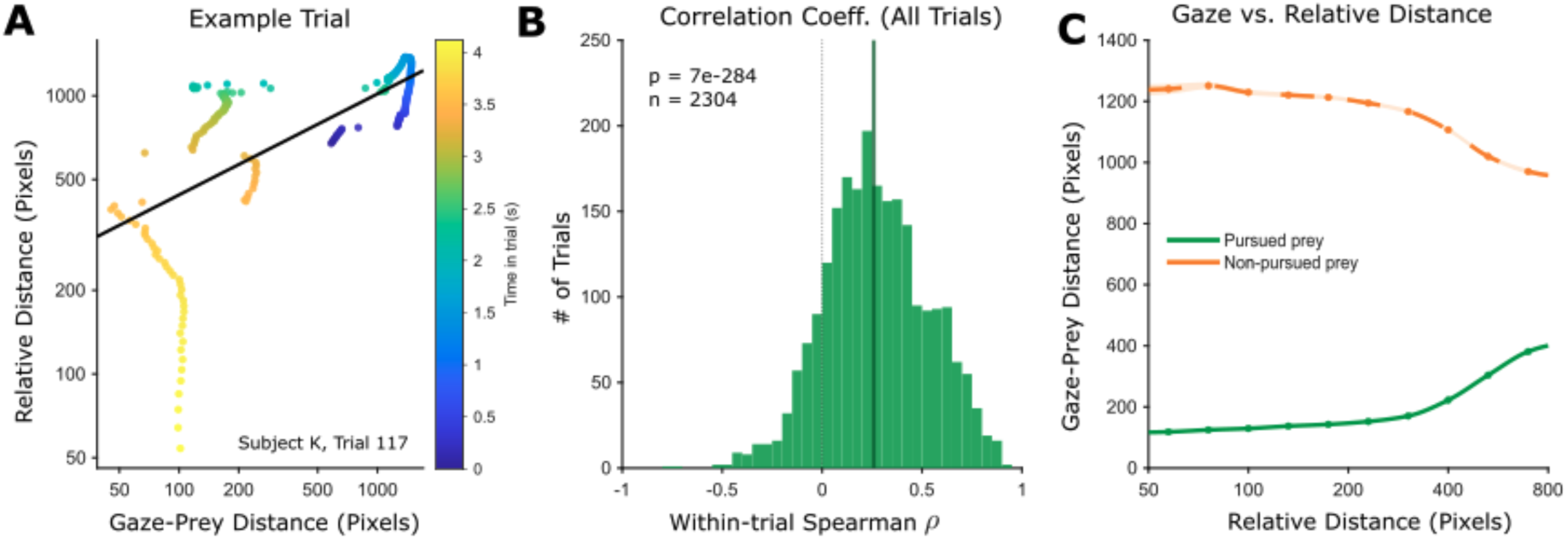
Gaze and Relative Distance are Correlated. **a)** shows an example trial for gaze-prey distance and the relative distance. Each dot represents a frame within the trial, and the color bar indicates the time of the frame. b) shows all of the trial’s Spearman correlation coefficients, where the y-axis is the number of trials and the x-axis is the Spearman correlation coefficient. c) shows the average gaze-prey distance relative to distance in pixels. The yellow line indicates the non-pursued prey, while the green line indicates the pursued prey.

**Extended Data Fig. 6.**
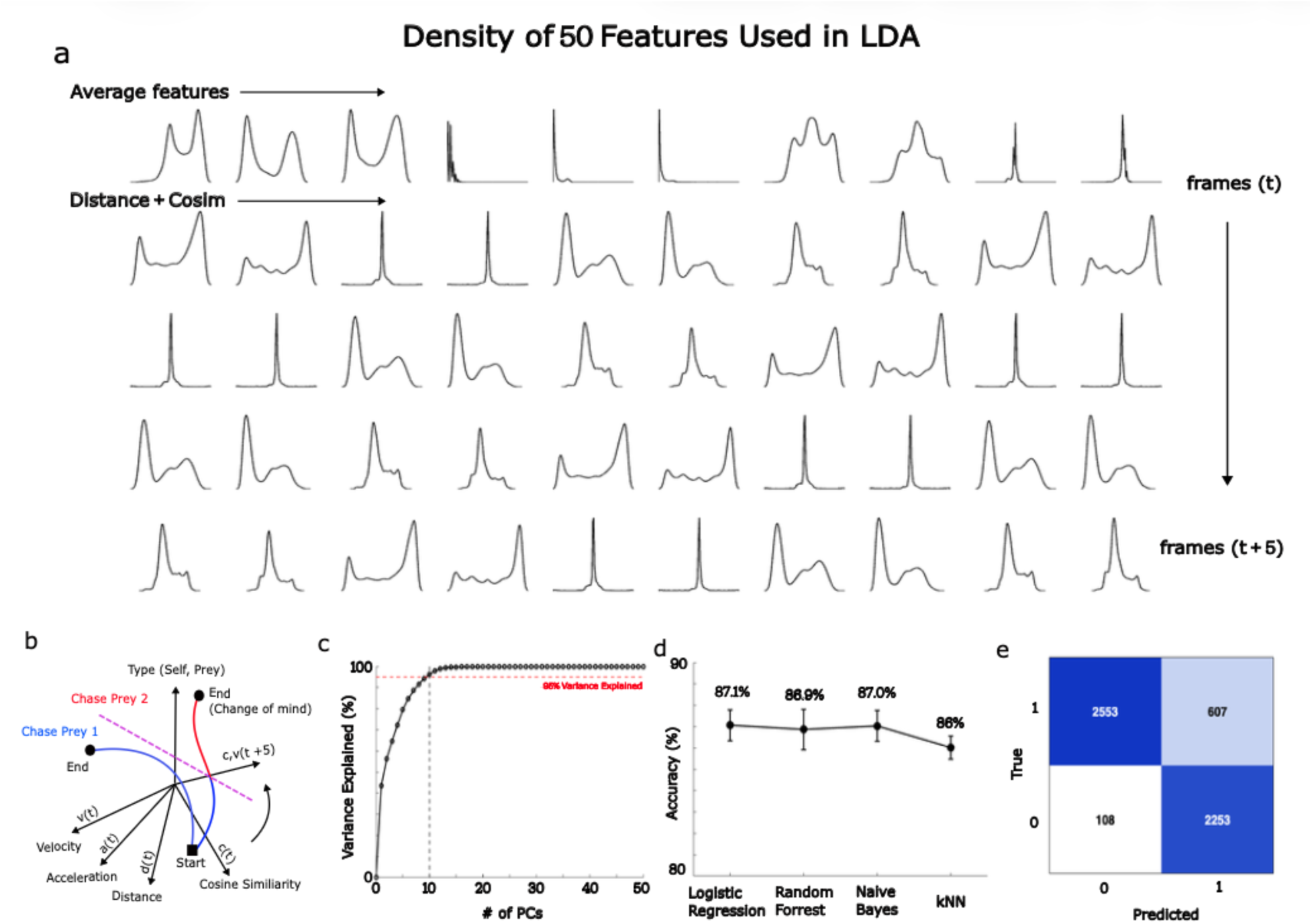
LDA Model for Detecting Change of Mind. **a)** The fifty features used to train the LDA model. The first line is the average of absolute features such as the agent’s velocity, the prey’s velocity, acceleration, and the difference in velocity. The next 40 features are relative distances and cosine similarities between the agent and the prey at time point “t” and over the next 5 frames (i.e., “t+5”). b) Schematics of the LDA model. If the trajectory crosses the line, it is a change-of-mind event. c) Scree plot of the PCA of the features. The black line indicates that approximately 10 PCs are needed to explain 95% of the variance. d) To test out the classifiers, we tried four different methods, including logistic regression, random forest, naive Bayes, and k-nearest neighbors algorithms. The best of these four models (logistic regression) was chosen. e) The error rate of each model after LDA classification. The model shows a preference for capturing prey 1 (class 0), with high precision (96%) but lower recall (81%). In contrast, prey 2 (class 1) has lower precision (79%) but higher recall (95%).

**Extended Data Fig. 7.**
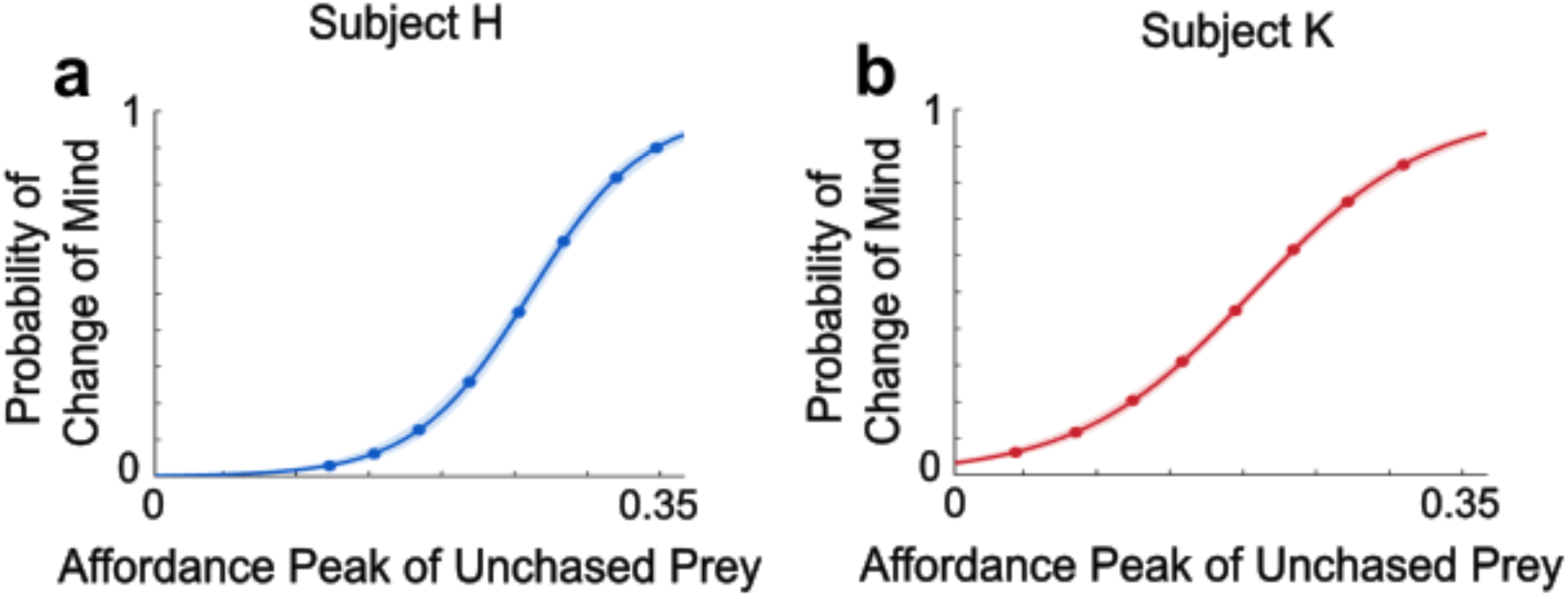
Psychometric Curve of Change of Mind Based on the Affordance Peak of Unchased Prey. **a)** Shows the psychometric curve of subject H. The y-axis is the probability of change of mind, while the x-axis is the peak affordance value of the original unchased prey. b) The same but for subject K.

**Extended Data Fig. 8.**
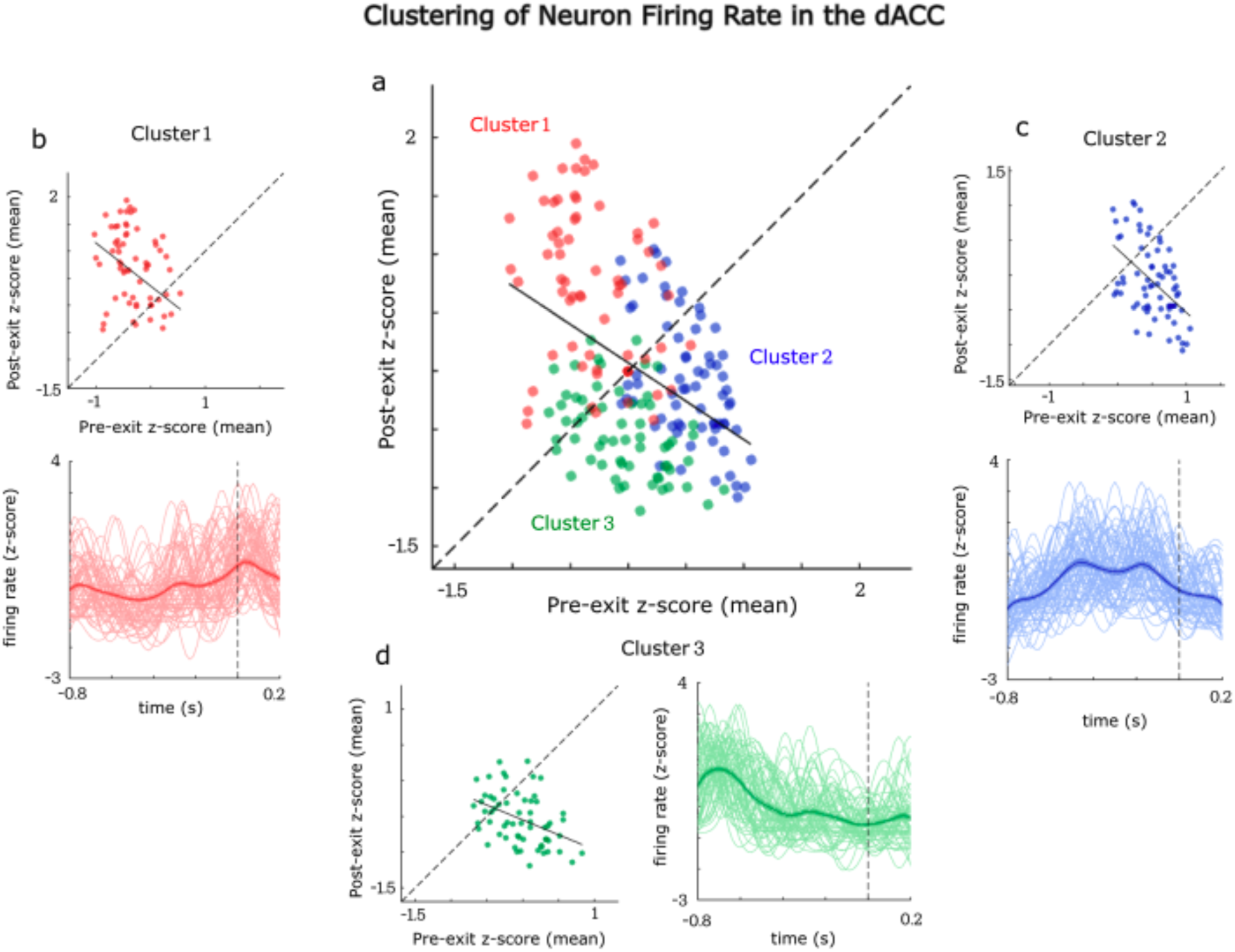
Firing Rate of Cluster of Neurons in the dACC. **a)** k-nearest neighbor plot for all the neurons recorded in the dACC (n = 218). The red represents cluster 1, the blue represents cluster 2, and the green represents cluster 3. b) Cluster 1, as shown in the scatter plot, has a higher firing rate after the exit compared to the pre-exit conditions. Exit, in this case, means the model no longer becomes confident that the subject is chasing the original prey. c) Cluster 2 shows ramping of the firing rate before the exit. d) Cluster 3 shows a transient during the start of the trials and inhibits near the exit. The cluster of peak mean firing rate forms before the exit.

## Supplementary Tables

**Supplementary Table 1.**
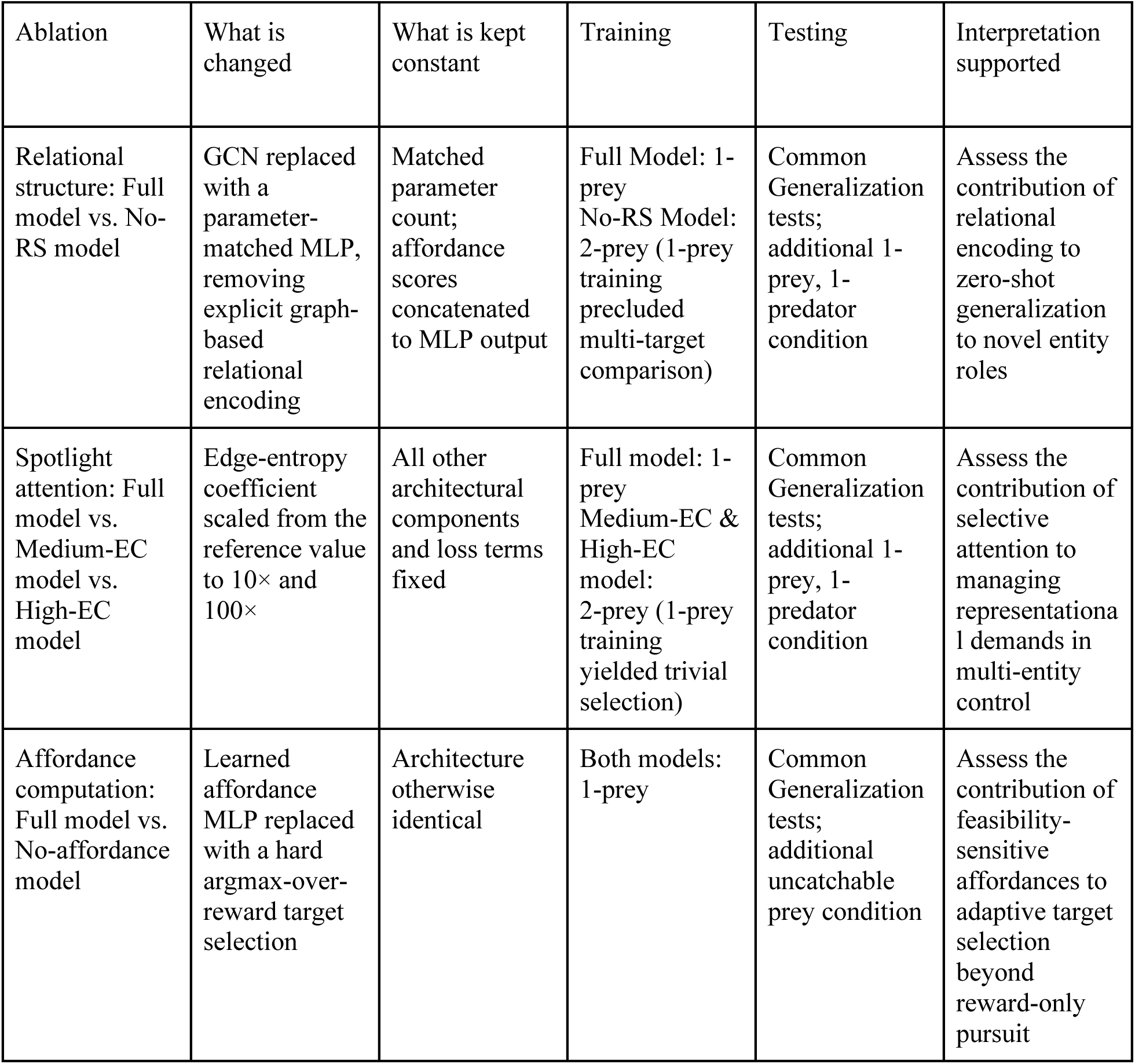
Summary of model comparisons and ablation analyses. All models were trained using the same environment. 1280 x 720 pixels, prey speed range [4, 8] pixels/timestep, and a nominal surface-friction parameter (agent = 0.1, prey = 0.15). All models were evaluated using a common generalization test that included expanded arena sizes, variation in prey speed range, surface friction, and increased prey numbers.

| Ablation | What is changed | What is kept constant | Training | Testing | Interpretation supported |
|---|---|---|---|---|---|
| Relational structure: Full model vs. No-RS model | GCN replaced with a parameter-matched MLP, removing explicit graph-based relational encoding | Matched parameter count; affordance scores concatenated to MLP output | Full Model: 1-prey<br>No-RS Model: 2-prey (1-prey training precluded multi-target comparison) | Common Generalization tests; additional 1-prey, 1-predator condition | Assess the contribution of relational encoding to zero-shot generalization to novel entity roles |
| Spotlight attention: Full model vs. Medium-EC model vs. High-EC model | Edge-entropy coefficient scaled from the reference value to 10× and 100× | All other architectural components and loss terms fixed | Full model: 1-prey<br>Medium-EC & High-EC model:<br>2-prey (1-prey training yielded trivial selection) | Common Generalization tests; additional 1-prey, 1-predator condition | Assess the contribution of selective attention to managing representational demands in multi-entity control |
| Affordance computation: Full model vs. No-affordance model | Learned affordance MLP replaced with a hard argmax-over-reward target selection | Architecture otherwise identical | Both models: 1-prey | Common Generalization tests; additional uncatchable prey condition | Assess the contribution of feasibility-sensitive affordances to adaptive target selection beyond reward-only pursuit |

**Supplementary Table 2.** Model names and definitions. Standardized model names, previously used terms, and defining differences.

| Unified label | Prior terms used in the manuscript | Definition |
|---|---|---|
| Full model | base model; affordance-based model; GCN-RL; with-RS model | Complete architecture (affordance module, GCN, RNN, PPO actor–critic), edge-entropy coefficient *c<sub>edge</sub>* = 1× (reference value) |
| No-RS model | without-RS model; | GCN replaced with a parameter-matched MLP |
| No-affordance model | without-affordance model; reward-only model; reward-based model; | Learned affordance MLP replaced with a hard argmax-over-reward target selection |
| High-EC model | control model | Edge-entropy coefficient *c<sub>edge</sub>* scaled to 100× the reference value, producing near-uniform attention |
| Medium-EC model | medium model; medium-temperature model | Edge-entropy coefficient *c<sub>edge</sub>* scaled to 10× the reference value |

## Methods

### Animal procedure

We utilized a previously published behavioral dataset collected from two male rhesus macaques to benchmark the agent’s performance^32^. The subjects were aged 9 years (subject K) and 10 years (subject H). The original data were collected under an animal-use protocol held by co-author Benjamin Hayden. All procedures adhered to the guidelines of the Public Health Service’s Guide for the Care and Use of Laboratory Animals and were approved by the Institutional Animal Care and Use Committees (IACUCs) at both the University of Rochester and the University of Minnesota.

### Behavioral task

We adopted a primate-inspired pursuit task to evaluate decision-making in dynamic environments^32^. In each trial, an agent controlled an avatar that pursued a moving prey within a bounded two-dimensional arena (training: 1280 × 720; testing: 1920 × 1080 pixels). The prey followed an adaptive avoidance policy that balanced keeping distance from the agent with staying within the arena. At each time step, it evaluated 15 candidate future positions and chose the one minimizing a cost function combining an environment-dependent term (including a bias toward the arena center) and an agent-dependent term. The agent term was a sigmoidal function of agent–prey distance, rising nonlinearly as the agent approached. If the best position violated boundaries, the prey selected the next-best feasible option.

### Experimental apparatus and controls

Subjects controlled the avatar using a modified joystick (Logitech Extreme Pro 3D) capped with a 3-cm sphere. Analog input was sampled and mapped to avatar motion using custom MATLAB code. Movement obeyed a kinematic constraint: maximum displacement was capped at 23 pixels per frame (16.67 ms). We used the same constraint to define the artificial agent’s maximum velocity, ensuring kinematic equivalence between biological and computational control.

### Training environment for agent

The RL objective used a composite reward balancing efficiency and value. On capture, the agent received a scalar reward (12–108) proportional to the prey’s maximum speed, incentivizing pursuit of faster, higher-value prey. Each time step incurred a small time penalty (−0.0001) to encourage rapid interception, and failure to capture within 500 timesteps triggered a terminal penalty.

### Artificial networks

We developed a modular deep RL architecture comprising four sequential components: affordance computation, attention-weighted relational structure (graph convolutional network; GCN^34,35^), temporal integration (gated recurrent unit; GRU^36^), and decision-making (PPO actor–critic^37^). The model processes observations through an affordance-based filtering stage, constructs relational representations using a continuously weighted GCN, integrates these representations over time with the GRU, and maps the resulting state to continuous actions via PPO. At each time step *t*, the agent observed an object-centric state matrix *X_t_* ∈ R*^N^*^×^*^F^*, with *N* = 6 entities and *F* = 5 features, where the first row represented the agent, and the remaining five rows represented potential prey. To prevent position-dependent heuristics tied to input ordering, prey rows were randomly permuted at every step. Each row contained a normalized feature vector *x_t_*_,*i*_ (scaled to [0, 1]) comprising the entity’s 2D position, 2D velocity, and a scalar encoding its reward identity (equivalently, its color), whose value was determined by the prey’s maximum speed. During 1-prey trials, unused positions were zero-padded and masked during attention computation.

The affordance module computed the relative task relevance of each observable entity at each time step. Each entity was processed independently by a shared multilayer perceptron (MLP), consisting of two fully connected layers with 64 hidden units and ReLU activation, which first computed a per-target score from its position, velocity, and reward magnitude,

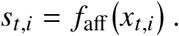

Applying the same MLP parameters to all entities promotes generalization across targets. To impose a competitive relevance constraint, these scores were converted into GCN edge weights, *α_t_*_,*i*_, by applying a masked softmax,

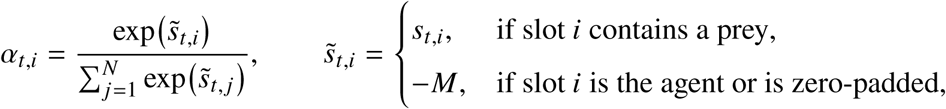

where *M* is a large positive constant (*M* =10^9^). The agent’s own node and any zero-padded slot were assigned a large negative score prior to the softmax, driving their weight to effectively zero. Consequently, two targets with equal reward generally receive different edge weights if they differ in position or velocity. These weights defined symmetric connections between the agent and each prey, forming the affordance-derived adjacency matrix *A_t_* ∈ R*^N^*^×^*^N^* that determines the strength of information flow between nodes.

The GCN first applied a shared linear projection to map each entity’s state features to a 64-dimensional representation,

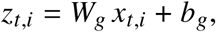

where the projection weights *W_g_* were initialized using Glorot uniform initialization^54^ and the bias *b_g_* was initialized to zero. It then computed each node’s updated representation as a weighted sum of its neighbors’ projected features, using the affordance-derived edge weights,

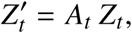

where the rows of *Z_t_* are the projected embeddings *z_t_*_,*i*_. This formulation implements a weighted message-passing mechanism in which each entity’s influence on the relational representation is scaled by its affordance weight: targets assigned larger edge weights contributed more strongly to the agent’s updated representation, while targets assigned near-zero weights contributed correspondingly little. The resulting per-entity representations were flattened and concatenated into a single vector *v_t_*, a compact representation of the current spatial scene weighted by affordance-driven attention, and passed to the recurrent module for temporal integration. This module was a single-layer GRU with a 512-dimensional hidden state ℎ*_t_*, updated iteratively from the current input *v_t_* and the previous hidden state ℎ*_t_*_−1_,

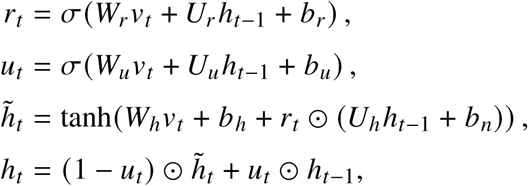

where *r_t_* and *u_t_* are the reset and update gates, *b_n_* is a bias on the recurrent candidate term, *σ* is the logistic sigmoid and ⊙ denotes element-wise multiplication. The GRU hidden state served as the latent state and sole input to the PPO actor–critic module. The actor produced a continuous 2D action vector constrained to the unit disk (see PPO module), matching the analog joystick control used in the primate experiments and ensuring kinematic equivalence between biological and artificial agents (Fig. 1b).

Policy learning used PPO with Adam^55^ (learning rate 1 × 10^−5^, *β*_1_ = 0.9, *β*_2_ = 0.999, *ε*_Adam_ = 1 × 10^−8^), clipping *ε* = 0.2, discount *γ* = 0.99, and generalized advantage estimation (GAE)^56^ *λ* = 0.95. We included entropy regularization (0.01), weighted the value loss by 0.5, and applied auxiliary entropy regularization to the affordance-derived adjacency matrix with a coefficient of 1.0 (see PPO module). Trajectories were collected from 64 parallel environments in 128-step rollouts (8,192 transitions per update); each update used one optimization epoch over 32 minibatches (minibatch size 256). To stabilize learning, we applied global gradient-norm clipping with a maximum norm of 0.5 and normalized advantages within each update batch. Agents were trained for ≥ 250,000,000 environmental steps. For ablations, training continued until a predefined success criterion was met, with all hyperparameters held constant across conditions.

### PPO module

The PPO module maps the recurrent state to actions and value estimates and optimizes all network parameters end-to-end. Policy learning and value estimation are decoupled through separate actor and critic networks. The actor network takes the current state ℎ*_t_* as input and consists of a single hidden layer with 128 units and ReLU activation, followed by a linear output layer that produces the mean action vector *μ_t_* for the continuous action space. Network weights are initialized orthogonally^57^, with a gain of 2.0 for the hidden layer and 0.01 for the output layer. The policy is parameterized as a multivariate Gaussian distribution with diagonal covariance,

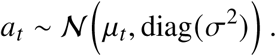

Where the log standard deviation log *σ* is a learned, state-independent parameter shared across time steps. The critic network receives the same recurrent hidden state and estimates a scalar state-value function *V* (ℎ*_t_*). It consists of a single hidden layer with 128 units and tanh activation. The critic network is initialized orthogonally, with a gain of 2.0 for the hidden layer and 1.0 for the output layer. Sampled actions were first clipped component-wise to [−1, 1] by the environment wrapper. During the movement update, clipped vectors with Euclidean norm greater than 1 were divided by their norm; vectors within the unit disk were unchanged. This radial rescaling preserved the direction of the clipped vector. The resulting control vector was then used to compute the target velocity. All parameters are optimized jointly using PPO with the Adam optimizer by minimizing the objective,

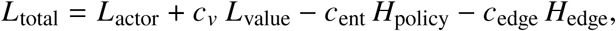

where *c_v_* = 0.5, *c*_ent_ = 0.01, and *c*_edge_ weight the value loss, policy entropy, and edge-entropy terms, respectively.

*L*_actor_ represents the negative clipped surrogate policy loss, defined as:

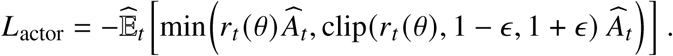

Here, *r_t_* (*θ*) = *π_θ_* (*a_t_* | ℎ*_t_*)/*π_θ_*_old_ (*a_t_* | ℎ*_t_*) denotes the importance-sampling ratio between the probability density assigned to the sample action under the updated policy (*π_θ_*) and the assigned under the old policy (*π_θ_*_old_ (*a_t_* | ℎ*_t_*)), which generated the rollout. *Â_t_* is the generalized advantage estimate computed from the rollout, and *ε* = 0.2 is the clipping threshold.

*L*_value_ is the clipped value function loss, ensuring the value updates do not deviate excessively from the previous predictions:

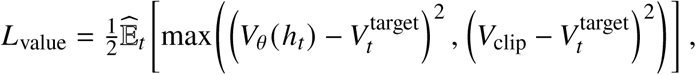

where *V_θ_* (ℎ*_t_*) is the critic’s current value estimate,

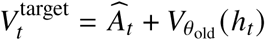

is the value regression target, and

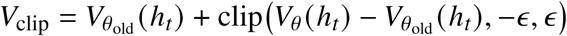

is a clipped value estimate that limits how much the value prediction can change in a single update, following the same clipping threshold *ε* used in *L*_actor_. *H*_policy_ denotes the standard policy entropy. *H*_edge_ denotes an auxiliary entropy regularizer computed from the affordance-derived adjacency matrix used for relational aggregation.

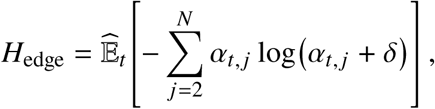

where *α_t, j_* denotes the validity-masked edge weights assigned to the *j*-th of the *N* − 1 potential target slots at time *t*. The agent’s own node is excluded from this distribution before the softmax, while any zero-padded target slot remains in the distribution but is assigned a large negative score before the softmax, giving its weight effectively zero and making no meaningful contribution to *H*_edge_. *δ* is a small constant (10^−8^) for numerical stability. We fixed the edge-entropy coefficient at its reference value, *c*_edge_ = 1.0, for the Full model. This regularization discourages premature collapse of the affordance-derived relational structure and promotes diffuse relational interaction during learning.

### Ablation of the spotlight attention mechanism

To examine the contribution of the spotlight attention mechanism, we implemented ablation models in which the selective capacity was systematically attenuated. We modified the distribution of affordance-derived attention weights to approximate a divided-attention regime. This was implemented by adding an edge-entropy regularization term to the PPO objective function. This term penalizes highly peaked attention weights, encouraging the model to distribute its focus more evenly. These models use the identical loss formulation defined in the PPO module. The only change relative to the Full model is the edge-entropy coefficient *c*_edge_, which we increased to 10x and 100x its reference value (i.e., *c*_edge_ = 10 and *c*_edge_ = 100) for the Medium-EC and High-EC models, respectively. Because *L*_total_ includes −*c*_edge_*H*_edge_, increasing *c*_edge_ more strongly encourages higher-entropy attention distribution during training, while all other loss terms and coefficients remain unchanged. By scaling the edge entropy coefficient, we control the degree of focus during target selection. Under a high edge-coefficient condition, this formulation produces a network that learns a uniform attention distribution across entities, reducing sensitivity to differences in affordance scores. Under a medium edge-coefficient condition, the distribution became less sharp, leading to partial dispersion of attention across entities. Unlike the Full model, which was trained in a 1-prey environment, this ablation model was trained in a 2-prey environment. This design choice was motivated by a preliminary observation that when trained on a 1-prey, the model converged on trivial selective behavior regardless of entropy strength, because the absence of competing targets eliminated the need for attentional allocation. Training with 2-prey introduced a minimal competitive context, in which the high edge coefficient setting led to equally distributed attention rather than a focus on a single target.

### Ablation of relational structure

To assess the role of explicit relational structure in representing interactions among multiple entities, we implemented an ablation model that removed graph-based relational modeling. In this variant, the GNN module was replaced with a standard MLP, eliminating inductive biases that explicitly encode relationships between the agent and surrounding entities. To ensure that any observed performance differences were attributable to the absence of relational structure rather than to differences in representational capacity, the substituted MLP was designed to match the original GCN in total learnable parameters. The MLP consisted of a fully connected layer with 64 hidden units, followed by a ReLU activation. To prevent the loss of affordance information in this ablation, the affordance scores computed for each entity were concatenated to the MLP input before being passed to the subsequent temporal processing module. This design ensured that affordance signals remained available to the policy, isolating the effect of removing relational structure while preserving information about target relevance. The model was trained in an environment containing exactly two targets. Preliminary experiments showed that non-relational models trained with a single target failed catastrophically when evaluated in multi-target settings, precluding meaningful assessment of reward-based decision-making, such as selecting higher-value targets in competitive environments. Training with two targets provided a minimal functional baseline, enabling valid comparison of pursuit strategies in the absence of explicit relational structure.

### Ablation of affordance computation

To assess the role of the state-dependent affordance computation module, we implemented an ablation model that removed the learned affordance computation. In the Full model, affordance scores reflect not only the intrinsic value of a target but also its feasibility relative to the agent’s current state. This ablation was designed to examine the behavioral consequences of eliminating this adaptive evaluation process. In this No-affordance model, the learnable multi-layer perceptron that computes affordance scores was removed and replaced with a deterministic heuristic. Instead of deriving attention weights from learned affordance values, the model selected targets solely based on their predefined intrinsic reward magnitudes. Consequently, attention was assigned using an argmax operation over the reward values of all visible entities:

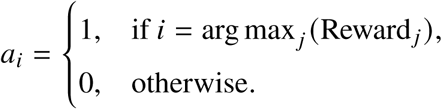

This substitution forced the agent to adopt a reward-based strategy, in which it always pursued the target with the highest reward. As a result, the agent disregarded contextual factors such as spatial distance, relative velocity, and time constraints that influence whether a target is practically attainable. By contrasting this heuristic policy with the Full model under the same zero-shot evaluation without additional training, this ablation examined the contribution of learned affordance computation, comparing pursuit based on reward magnitude with pursuit guided by state-dependent affordance score. The target-selection rule in the No-affordance model remains fixed, although the downstream policy is trained using PPO.

### Generalization protocols and evaluation metrics

To evaluate the robustness and adaptability of the learned policies, we subjected all agents to a series of zero-shot generalization tests. These evaluations were conducted in environments that deviated substantially from the training distribution, probing the agents’ ability to cope with unseen physical dynamics, altered kinematic demands, and increased task complexity without additional learning. During training, all models were exposed to a fixed environmental configuration with a resolution of 1280 × 720, prey speed sampled uniformly from [4, 8] pixels per frame, and a nominal surface-friction parameter (agent = 0.1, prey = 0.15). Generalization performance was assessed across four categories of environmental perturbations.

First, to test spatial generalization, agents were evaluated in expanded arenas with resolutions of 1920 × 1080 (corresponding to the primate experimental setup) and 2560 × 1440. Second, to assess robustness to kinematic shifts, prey speed ranges were increased to [7, 11] and [10, 14] pixels per timestep, exceeding the range encountered during training and requiring more precise predictive control. Third, to examine sensitivity to changes in control dynamics, surface friction coefficients were systematically altered. Relative to the nominal condition, we introduced reduced-friction environments (*υ* − 0.1) and increased-friction environments (*υ* + 0.1), perturbing the mapping between control actions and resulting motion. Finally, to evaluate performance under increased task complexity and attentional demands, agents were tested in multi-prey environments with three to five targets. Training conditions for prey number differed across models to ensure meaningful evaluation. The Full model and the No-affordance model were trained with a 1-prey. In contrast, the No-RS model and the Medium-EC and High-EC models were trained with 2 prey; training with 1 prey in these cases failed to elicit competitive selection behavior, thereby masking the functional role of the removed components.

In addition, we designed a generalization test to specifically probe the role of affordance computation under extreme kinematic conditions. This evaluation was motivated by the observation that models without affordance computation tend to pursue the highest-value target regardless of its feasibility. To test whether the Full model could adaptively abandon an unattainable target, we applied this condition selectively to the Full model and to the ablation model without affordance computation. In this test, the maximum speed of the highest-value prey was increased from 14 to 30 pixels per timestep, exceeding the agent’s effective interception capability within the fixed time limit. Successful performance in this regime, therefore, required the agent to evaluate the actionability of available targets and to prioritize lower-value but attainable prey over an unreachable high-value option. This condition tested whether the learned policy could integrate value information with feasibility constraints in a zero-shot manner, without additional training.

Finally, we introduced a mixed-valence test scenario to assess whether agents could distinguish between rewarding targets and aversive threats that had never been encountered during training. This condition tested the learned policy’s ability to integrate relational and reward-based signals when encountering qualitatively novel entities. In this scenario, the environment contained one prey species and one predator species. The prey followed the same dynamics as in the standard task, whereas the predator was programmed to pursue the agent at a fixed speed of 4 pixels per timestep, which was slower than the agent’s speed. If the agent was caught by the predator, the episode terminated immediately, and a fixed penalty of −10 was received. The agent needs to avoid the predator while continuing to pursue the target.

The predator was never present during training. To encode its aversive nature at test time, the predator’s intrinsic reward value in the state matrix was set to −0.001. In addition, unlike standard targets whose attention weights were derived from the affordance module, the predator’s affordance weight was manually set to −0.1 within the graph representation. This configuration allowed us to examine whether the learned policy could respond appropriately to an unseen threat solely based on relational and reward-based cues, without additional training. Performance in this mixed-valence setting was compared between the Full model and ablation variants lacking spotlight attention or relational structure, enabling assessment of how these components contributed to generalization in the presence of novel negative entities. To further examine the contribution of relational aggregation in the Full and No-RS model comparison, we performed an edge-weight swap analysis on the trained Full model. At each evaluation step, the prey and predator edge weights were exchanged in both directions of the agent-entity connections before graph aggregation, without rearranging node features or modifying learned parameters. No additional training was performed.

Performance was quantified using two complementary metrics. Capture success rate measured the proportion of episodes in which the agent successfully intercepted any prey within the time limit, providing an index of basic sensorimotor competence. The success rate of higher value measured the proportion of episodes in which the agent intercepted the highest-value prey among multiple available targets, thereby assessing selective attention and reward-based decision-making under competition. Success rates of the Full model were compared with those of the subjects and of the No-RS model using chi-squared tests; success rates across seeds were compared between models using Welch’s *t*-tests; and the edge-weight swap was compared with the intact Full model using a two-tailed paired *t*-test across seeds. The test used for each comparison is reported with the corresponding result.

### Behavioral affordance

We developed a method to quantify the moment-to-moment action relevance of each prey. This temporal discount affordance model assigns an affordance value to each prey at each frame to explain the prey’s immediate pursuability. First, we computed the time-to-contact (TTC) function between the self and each prey as a function of time, where *D* is denoted as the Euclidean distance between self and prey, and *V* denotes the respective speeds at a specific time (*t*). TTC can only be defined when *v*_self_ > *v*_prey_, reflecting that capture is not kinematically feasible under the condition where *v*_self_ < *v*_prey_.

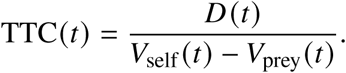

TTC was expressed in seconds; when *V*_self_ ≤ *V*_prey_, or when TTC exceeded the 20-s trial limit, TTC was set to 20 s. The TTC value is then transformed into an affordance value via an exponential temporal discounting function:

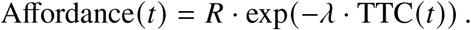

*R* represents the theoretical reward associated with the specific prey, and (*λ* = 0.4) is a temporal discounting parameter controlling the sensitivity to delay. Theoretically, this equation assigns higher affordance to prey that are closer, relatively slower, and higher-reward, consistent with the normative principles for chasing. Notably, the affordance measure is computed continuously over time and does not assume explicit planning or value learning. Instead, it captures the instantaneous action potential of each prey given the current kinematic state, providing a behaviorally grounded signal suitable for comparison with both choice behavior and neural activity. Unlike conventional value presentation, this affordance signal is inherently state-dependent and dynamically evolves with the kinematic feasibility of capture, rather than reflecting a stationary expected reward.

### Cosine similarity analysis

Similarity between subject and agent affordance matrices was quantified using cosine similarity:

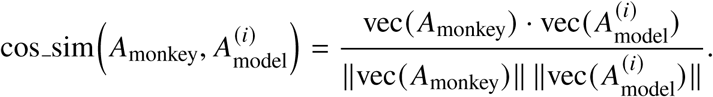

Uncertainty in the mean cosine similarity was estimated by bootstrap resampling across seeds (10,000 resamples). Significance was tested using a one-sided sign-flip randomization test across seeds (200,000 permutations) against the null hypothesis of zero mean similarity.

### Behavioral similarity analysis

Beyond the similarity of affordance structure, we quantified the fidelity of the artificial agent’s moment-to-moment decision-making relative to the biological subjects using a state-initialized trajectory-forecasting test. This test examined whether the agent, when initialized in the exact kinematic state observed for a primate subject, generated pursuit trajectories that aligned with the biological ground truth. From the behavioral dataset, we randomly sampled trials for each subject and task and, within each selected trial, randomly chose a reference time step *t*_ref_ as the initialization point. The agent was initialized with the precise kinematic states of both the subject’s avatar and the prey at *t*_ref_, including position *p* and velocity *v*. From this synchronized state, the agent interacted autonomously with the environment over a 10-time-step prediction horizon, producing a predicted trajectory. Behavioral deviation was quantified as the root-mean-square error (RMSE) between the agent’s predicted position *p*^ and the subject’s observed position *p*; RMSE was first computed separately for each time step within the horizon and then averaged across time steps to yield a single trajectory-level error for each sample, with lower values indicating closer alignment with the biological trajectories. To compare behavioral fidelity across agents, we fit a linear mixed-effects model with agent type as a fixed effect and random effects accounting for repeated measurements across trials and subjects, thereby controlling for inter-subject variability and within-subject correlations.

### Detecting change of mind using a linear discriminant classifier

To infer which prey the subject was pursuing at each moment, we built a supervised, frame-wise classifier and used it to detect change-of-mind events in 2-prey trials. For each video frame, we computed a 50-dimensional feature vector capturing the subject–prey interaction dynamics: 10 instantaneous kinematic variables (the speeds and acceleration magnitudes of the avatar and each prey, and the avatar–prey differences in speed and in acceleration) and 40 relational terms evaluated at five future offsets (*t* + 1 to *t* + 5 frames), namely the cosine similarity between the avatar’s velocity and each prey’s velocity, the distance between the avatar and each prey, and the frame-to-frame change in each. To construct a training set with high-confidence labels, we extracted the final 60 frames of each successful 2-prey trial, in which the captured prey was known, assuming that behavior near capture reflects a committed pursuit. Labels were assigned based on which prey’s reward value was ultimately obtained (prey 1 vs prey 2). Features were z-scored using the per-feature mean and standard deviation computed across the training set and reduced with PCA, retaining 20 components; the top 10 principal components (explaining approximately 95% of variance) were used to train a linear discriminant analysis (LDA) classifier, which projected the data onto a single discriminant axis that maximally separated prey-1-captured from prey-2-captured trials. This axis yielded a one-dimensional decision variable indexing relative evidence for pursuing prey 1 versus prey 2. To convert this decision variable into calibrated probabilities, we fit a logistic regression mapping from the LDA output to *P*(pursue prey 2) (selected from a small candidate set of link-function models). We then applied the model to every frame of all 2-prey trials to obtain a continuous, moment-by-moment confidence trace of pursuit probability. A trial was classified as a change of mind only if it satisfied three criteria designed to ensure interpretable, sustained commitment: (i) an initial commitment phase, occurring before any commitment toward the finally captured prey, in which the posterior lay within 0.15 of the bound for the prey that was not ultimately captured (*P* < 0.15 for prey 1; *P* > 0.85 for prey 2) for at least 30 consecutive frames (0.5 s); (ii) a subsequent transition in which the posterior crossed into the bound of the alternative prey, with the initial commitment ending before the first frame of this new commitment (avoiding “grey-zone” fluctuations); and (iii) the initial committed prey was not the final captured prey (non-final choice preceding the final choice). For each change-of-mind trial, we further defined the behavioral exit event as the first frame at which the LDA confidence trace crossed a threshold of |LDA| = 0.5 away from the initial commitment direction, where the initial commitment direction was determined from the first 25% of the trial or the first 60 frames, whichever was shorter. These inferred pursuit states, change-of-mind timestamps, and exit events were used in all subsequent behavioral and neural analyses.

### Affordance dynamics of change of mind

To quantify the behavioral correlates of change-of-mind events, we analyzed both static and dynamic properties of the behavioral affordance signals defined above (see Behavioral affordance), which were computed continuously for each prey throughout the trial. For each trial, we computed five static affordance statistics, including the mean, standard deviation, and pairwise differences between competing prey, together with five dynamic features capturing the temporal structure of affordance evolution, including momentum, peak rate of change, and trend strength. These ten affordance-derived variables were used as predictors in a linear regression model to classify whether a given trial contained a change-of-mind event. Because affordance was tracked continuously, we could also identify the time points at which the affordance values of the two prey crossed, and from these crossings we extracted the timing of affordance crossover events. LDA was then applied to characterize the temporal structure of three event types: affordance crossover, exit, and entrance (the first frame of commitment to the newly pursued prey). Finally, to assess how much temporal information was required for accurate decoding, we systematically varied the temporal window used as model input and quantified decoding performance as a function of the available temporal context.

### Neural recordings and preprocessing

Single-unit activity was recorded from the dorsal anterior cingulate cortex (dACC) across 42 sessions (5 sessions from subject H and 37 sessions from subject K) and sampled at 60 Hz. For condition-averaged analyses (PCA, alignment, and Procrustes), firing rates were extracted from a 60-frame (1 s) window beginning at frame 31 of each trial, and each neuron’s condition-averaged peri-stimulus time histogram (PSTH) was smoothed with a Gaussian kernel (*σ* = 3 bins, 31-bin window). For single-trial analyses (change-of-mind decoding), a separate Gaussian kernel (*σ* = 10 bins, 61-bin window) was used. For each neuron, the 1-prey (P1) and 2-prey (P2) condition-averaged PSTHs were then combined, and the global mean *μ_n_* and range were computed jointly across all conditions and time bins. Soft normalization was applied as

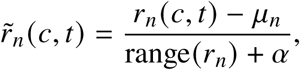

where *r_n_* (*c*, *t*) is the firing rate of neuron *n* in condition *c* at time bin *t* and *α* = 5 is a soft-normalization constant that prevents inflation of low-variance neurons. After soft normalization, the condition mean at each time bin was subtracted to remove shared temporal dynamics and isolate condition-dependent variance. These condition-mean-centered data formed the basis for all population-level analyses, including PCA, alignment, and Procrustes. To define affordance conditions, the behavioral affordance values of both prey were computed for each trial and summed as the area under the curve over the first second. The total affordance of each prey was compared across all sessions and binned by value; in 2-prey trials, binning used the affordance of the prey being pursued at the end of the first second.

### Dimensionality of population activity

PCA was performed on the population activity matrix *X* ∈ R^(^*^T^*^·^*^C^*^)×^*^N^*, with *T* = 60 time bins, *C* = 5 conditions, and *N* the total number of neurons, separately for the P1 and P2 conditions and for both the affordance and reward variables. The cumulative percentage of variance explained was plotted against the number of principal components (PCs), up to 50 PCs. To quantify uncertainty, we used a nonparametric bootstrap (100 iterations) in which the rows of the population matrix were resampled with replacement while preserving all *N* neurons, PCA was recomputed, and the cumulative variance explained was recorded; 95% confidence intervals were obtained as the 2.5th and 97.5th percentiles of the bootstrap distribution at each PC count. The number of PCs required to explain 90% of the variance (*k*_90_) was reported separately for P1 and P2, both as an absolute count and as a percentage of the total neuron count. To complement this measure with an estimate of effective dimensionality, we computed the participation ratio (PR)^58^ from the eigenvalue spectrum of the covariance matrix,

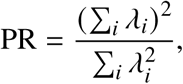

where *λ_i_* are the eigenvalues sorted in descending order. PR captures the number of dimensions that meaningfully contribute to population variance, with higher values indicating more distributed representations.

### Subspace alignment

To assess whether neural computations occupy a shared subspace across conditions, we computed the alignment index^59^ using the top 10 PCs. The alignment index quantifies the proportion of variance in one condition’s data that is explained by the principal-component subspace of another condition,

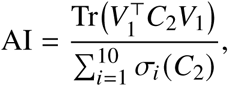

where *V*_1_ ∈ R*^N^*^×10^ contains the top 10 PCs from condition 1, *C*_2_ is the covariance matrix of condition 2, and *σ_i_* (*C*_2_) is its *i*-th largest eigenvalue. Statistical significance was assessed against a null distribution generated from tensor maximum-entropy (TME) surrogate data^60^, which preserve the marginal covariance structure along each tensor mode (time, condition, and neuron) while destroying higher-order dependencies between modes. To establish an upper bound on the alignment index achievable in the presence of recording noise, we used a split-half procedure. On each of 100 iterations (random seed = 42), the *N* neurons were randomly partitioned into two equal halves, with the same partition applied to both P1 and P2 data. The alignment index (P2 onto P1) was computed separately on each half and the two values were averaged, and the noise ceiling was reported as the mean across all 100 splits. This procedure estimates the alignment expected if P1 and P2 occupied the same subspace but were corrupted by independent noise in each half of the neurons.

### Cross-entity distance decoding

To test whether dACC represents the distance between the avatar and an entity in a format that generalizes across entities, we trained decoders on frames from one entity and tested them on frames from another. Units from both subjects with a mean firing rate of at least 0.5 Hz were pooled into a pseudo-population (186 units). Each unit’s activity was smoothed with a Gaussian kernel (*σ* = 2 frames). For each frame, the avatar–entity distance was assigned to one of 9 log-spaced bins between 50 and 800 pixels; frames closer than 50 pixels were assigned to the first bin and frames beyond 800 pixels were excluded. We tested three train–test pairs: (i) 2-prey trials in which prey 1 was pursued versus those in which prey 2 was pursued, labeled by the distance to the pursued prey; (ii) predator-free trials, labeled by the distance to the pursued prey, versus predator trials, labeled by the distance to the predator; and (iii) the same split as in (i), but labeled by the distance to the non-pursued prey. Because the non-pursued prey was rarely close to the avatar, we repeated analysis (iii) restricted to 500–1600 pixels (9 log-spaced bins), excluding frames outside this range. For each bin, 40 pseudo-trials were drawn by sampling each unit’s activity independently from frames of that bin, and this resampling was repeated 10 times. A Gaussian naive Bayes classifier with a uniform prior was trained on one side and tested on the other, in both directions, with small Gaussian noise (*σ* = 10^−6^) added to the training data to prevent singular variance estimates. Decoding accuracy was computed per bin as the fraction of test pseudo-trials assigned to the correct bin (chance = 1/9). Significance was assessed against 100 circular-shift surrogates, in which the distance labels of each trial were shifted by an independent random offset, preserving the autocorrelation of both firing and labels while breaking their alignment. For each bin and for the mean across bins, we report a *z*-score relative to the surrogate distribution and the corresponding one-sided *p*-value from a Gaussian fit to that distribution; per-bin *p*-values were corrected across bins with the Benjamini–Hochberg procedure.

### Gaze analysis

For subject K, for whom eye position was recorded, we computed in each 2-prey trial the distance between gaze and each prey and the distance between the avatar and each prey. Gaze samples that were missing or more than 200 pixels outside the screen were excluded, and trials shared across simultaneously recorded units were counted once (2,304 trials). The pursued prey was defined as the prey that came closest to the avatar during the trial. For each trial with at least 20 valid frames, we computed the Spearman correlation between avatar–prey distance and gaze–prey distance across frames, and tested whether the per-trial correlations differed from zero using a Wilcoxon signed-rank test.

### Single-neuron tuning

To characterize single-neuron tuning to task-relevant variables, we fit spline-based linear–nonlinear–Poisson (LNP) models^61^. For each neuron, we tested models incorporating reward (R), relative distance (D), and relative velocity (V), as well as their combinations (RD, RV, DV, and RDV). The linear stage computed a weighted sum of predictors, which was passed through an exponential nonlinearity to generate predicted firing rates. Model selection was performed using log-likelihood comparisons, and each neuron was classified according to the model or combination that best explained its activity; tuning was considered significant at *q* < 0.05 after Benjamini–Hochberg correction across neurons. Statistical significance of affordance tuning at the population level was assessed using a two-tailed binomial test.

### Neural correlates of change of mind

To examine neural activity surrounding changes of mind, neural data were aligned to the behavioral exit event defined above (see Detecting change of mind using a linear discriminant classifier). Prior to extraction, each neuron’s firing rate was z-scored using the session-wide mean and standard deviation, computed across all trials and time bins for that neuron, and then smoothed with a Gaussian kernel (*σ* = 10 frames, 61-frame window). We then extracted a window extending from 48 frames (0.8 s) before to 24 frames (0.4 s) after the exit event (73 frames in total at 60 Hz). To balance the comparison, non-CoM trials were randomly subsampled to match the number of CoM trials in each session (random seed = 42).

## Acknowledgment

This research was supported by the National Research Foundation of South Korea (RS-2023-00211018), (RS-2025-25431602), and the Institute of Information & Communications Technology Planning & Evaluation (IITP) with a grant funded by the Ministry of Science and ICT (MSIT) of the Republic of Korea in connection with the “Development of Human Cognition-Based AI Core Technology” (RS-2026-25518389). The authors declare no competing financial interests.

## Author contributions

Conceptualization: DHK, JSL, BYH, SBMY; Data Collection: SBMY; Analysis: DHK, JSL, SBMY; Writing—Original Draft: DHK, JSL, BYH, SBMY; Writing—Review and Editing: DHK, JSL, BYH, SBMY; Supervision: BYH, SBMY; Funding Acquisition: SBMY.

## Declaration of interest

The authors declare no competing interests. The funders had no role in study design, data analysis, or manuscript draft.

## Data availability

Data is available from the corresponding author upon reasonable request.

## Code availability

Code is available at https://github.com/Jungsugiee/Pursuit-Zeroshot

## Notes

### Competing Interest Statement

The authors have declared no competing interest.

### Summary of Updates

New version after revisions. The entire manuscript is updated.

